# Comprehensive in virio structure probing analysis of the influenza A virus identifies a functional RNA structure involved in replication and segment interactions

**DOI:** 10.1101/2020.03.05.975870

**Authors:** Naoki Takizawa, Koichi Higashi, Risa Karakida Kawaguchi, Yasuhiro Gotoh, Yutaka Suzuki, Tetsuya Hayashi, Ken Kurokawa

**Author notes:** Address correspondence to Naoki Takizawa.

## Abstract

The influenza A virus genome is segmented into eight viral RNAs (vRNA). Secondary structures on vRNA are thought to be involved in the viral proliferation process, such as intersegment interactions that are necessary for segment bundling. However, the functional RNA structure on vRNA is not well known because the secondary structure of vRNA in virion was partially unwound by binding viral non-specific RNA binding proteins in a sequence-independent manner. Here, we establish the global map of the vRNA secondary structure in virion using the combination of dimethyl sulfate (DMS)-seq and selective 2′-hydroxyl acylation analyzed by primer extension (SHAPE)-seq. By integrating DMS-seq and SHAPE-seq analyses with robust statistical analysis, we inferred quite a few bases paired regions including a pseudoknot structure on segment 5. Notably, when cells were infected with the recombinant virus which had mutations in the pseudoknot structure, the impairment of replication and packaging was observed on the other specific segment. Moreover, we analyzed the comprehensive intersegment RNA interactions in virion by ligation of interacting RNA followed by high-throughput sequencing (LIGR-seq). Our LIGR-seq analysis revealed that the intersegment interactions of the specific segment became less frequent and rearranged in the recombinant virus in concordance with the strength of genome packaging impairment. Our data provide evidence that the functional RNA structure motif on the influenza A virus genome can affect the efficiency of replication and segment bundling through the segment interactions.

## Introduction

The influenza A virus (IAV) genome consists of eight single-stranded negative-sense RNA segments (vRNA). One copy of each segment is packaged together into a single virus particle, and eight segments are organized in a conserved ‘7+1’ configuration in the virus particle [1, 2]. Segment reassortment is one of the driving forces for IAV evolution. Genetic reassortment between the human IAV and the avian/animal IAV can lead to the emergence of a new subtype of IAV, a candidate for a pandemic influenza strain. Each genome segment forms the viral ribonucleoprotein (vRNP) with the viral RNA polymerase and nucleoprotein (NP), a single-stranded RNA binding viral protein. In the previous study, the vRNPs in viral particles are revealed to form a double-helical structure with the polymerase at one end and a short loop at the other [3].

Previous studies have examined motif sequences required for efficient genome packaging and bundling [4]. The signal sequences for efficient genome packaging and bundling were initially found to be located in the coding regions at both ends of each segment, but they have also been found in the middle of coding regions. Mutations and deletions of some signal sequences resulted in impairment in the bundling of eight segments [5–10], and each segment was shown to have different importance for the viral genome bundling [11], suggesting that the bundling of the segmented genome is a hierarchical process. While the function of these signal sequences remains unknown, it is hypothesized to be involved in intersegment interactions for segment bundling. Regions responsible for the interactions between vRNAs have been identified *in vitro*, suggesting the existence of specific intersegment interaction networks necessary for genome packaging [12–15]. Direct contacts between the vRNPs have also been observed by electron tomographic analyses [16, 17]. Recent studies using a comprehensive high- throughput sequencing (HTS) approach demonstrated that a redundant and complex network of intersegment interactions found in the virion is essential for bundling the eight segments [18, 19]. As such, intersegment interactions are one of the important key factors to control precise genome bundling.

To form and regulate higher-order interactions including intersegment interactions, the importance of the RNA secondary structures has been widely recognized. In many RNA viruses, specific regions of the viral RNA genomes also act as cis-acting regulatory elements that mediate the virus propagation. These cis-acting RNA elements often form highly specialized structural motifs such as stem-loops or pseudoknots [20]. In IAV, the RNA structures at the promoter region, located at the 5′ and 3′ termini of the vRNA and their reformation at the promoter in transcription step, are elucidated by crystal structure and cryo-EM analyses [21–23]. The comprehensive analysis of specialized structural motifs coded on the IAV genome RNA other than promoter region was carried out by the RNA secondary structure predictions, identifying the conservation and enrichment of stem-loop structures on the IAV genome [24, 25]. Furthermore, the mutations that disrupt a predicted stem-loop and pseudoknot structure have been shown to reduce virus propagation [24–27]. Gavazzi et al. identified an *in vitro* direct interaction between segments 2 and 8 of an H5N2 avian IAV strain. They found that these two regions involved in the intersegment interaction may form stem-loop structures that can initiate the intersegment interaction by forming a kissing-loop complex [13]. These findings hypothesized that specialized structural motifs on the IAV genome RNA mediate intersegment interaction networks.

However, the fluctuation of vRNA structure in virion due to binding NP in a sequence-independent manner makes it hard to reveal the precise vRNA structure through *in silico* study. Recent studies by cross-linking immunoprecipitation (CLIP) analyses have revealed that NP does not bind vRNA uniformly, indicating that the secondary structures of vRNAs are partially unwound by binding NP. For that reason, some specific regions of the vRNP can form unpredictable secondary structures *in silico* [28, 29]. Dadonaite et al. revealed the secondary structures of the IAV genome in the virion using selective 2′-hydroxyl acylation analyzed by primer extension and mutational profiling (SHAPE-MaP). Their SHAPE-MaP profiles revealed that some vRNA secondary structures remain in the context of vRNP [19].

The comparative analysis of different HTS approaches showed that each HTS approach possesses the own detection bias for base reactivity [30]. Thus, to further understand the function of vRNA secondary structure, comprehensive and high- resolution information on the secondary structures of the IAV genome in virion needs to be created. In this study, we revealed the secondary structures of the IAV genome in the virion using multiple HTS technology and bioinformatics. We obtained a robust conformational map by combining two high-throughput and massive-scale sequencing techniques; dimethyl sulfate (DMS)-seq [31–33] and SHAPE-seq [34, 35], and multiple bioinformatical tools for calculating SHAPE reactivity; BUMHMM [36] and reactIDR [37]. DMS and NAI can modify different moieties of a single-stranded RNA [38], resulting in a different detectability depending on the sequence content. As a result, we identified a specialized structural motif on the vRNP and showed that the unwinding of this structural motif resulted in the impairment of replication. We further revealed the global intra- and intersegment interactions of the recombinant virus that had mutations in the structural motif region using ligation of interacting RNA followed by high-throughput sequencing (LIGR-seq) [39] and demonstrated that a part of intersegment interactions detected between multiple segments was rearranged in the recombinant virus. Our results suggest that the structural motif formed on vRNP is required for replication and segment interactions to preserve the genomic structures of the IAV.

## Results

### Identification of pseudoknot structure formed in the vRNP by high-throughput structure probing methods

To reveal the vRNA secondary structures by binding the viral proteins, we performed DMS-seq and SHAPE-seq for IAV genome RNA in three different conditions; the vRNA, vRNP, and virion. These methods are aimed to detect the RNA regions that are more accessible and likely to be attacked by the reagents. Thus, we can infer single-stranded and double-stranded regions at a single base resolution according to the reactivity scores. We utilized both DMS-seq and SHAPE-seq to uncover the whole landscape of the secondary structures of vRNA, which was highly complex with the viral proteins.

We carried out duplicate DMS-seq and SHAPE-seq experiments. The coverages were enough to calculate the reactivities except for the 3′ end of segments (Figure S1A). Reproducibility was evaluated by the drop-off rate of reverse transcriptase. The coefficient of determination of each duplicate experiment ranged from 0.26 to 0.93 (Figure S1B). To calculate a reliable score from these samples, we utilized robust statistical analyses. The probabilities of modifications for all nucleotides were calculated from the large-scale sequencing data using BUMHMM [36] and reactIDR [37]. reactIDR and BUMHMM output normalized probability that is an index of reactivity. The overall tendency of RNA structure was assessed for each segment using the violin plot of probabilities calculated by reactIDR (Figure 1A). As a result, the median of probability from the virion and the vRNP labeled with DMS was higher than that from the DMS- labeled vRNA (P-value < 2.2x10-16 by Kruskal-Wallis test), and those labeled with NAI was lower than that from the NAI-labeled vRNA in SHAPE-seq (P-value < 2.2x10-16 by Kruskal-Wallis test) (Figure 1A). Next, we compared probabilities of high-NP binding regions identified PAR-CLIP analysis [29] with that of the other regions (Tables S1 and S2). The probability of high-NP binding regions in vRNA was lower than that of the other regions. However, the probability of high-NP binding regions in vRNP and virion were higher than or comparable with those of the other regions. These results suggest that the overall secondary structure of vRNA is likely to be dissolved by forming in the vRNP.

**Figure 1.**
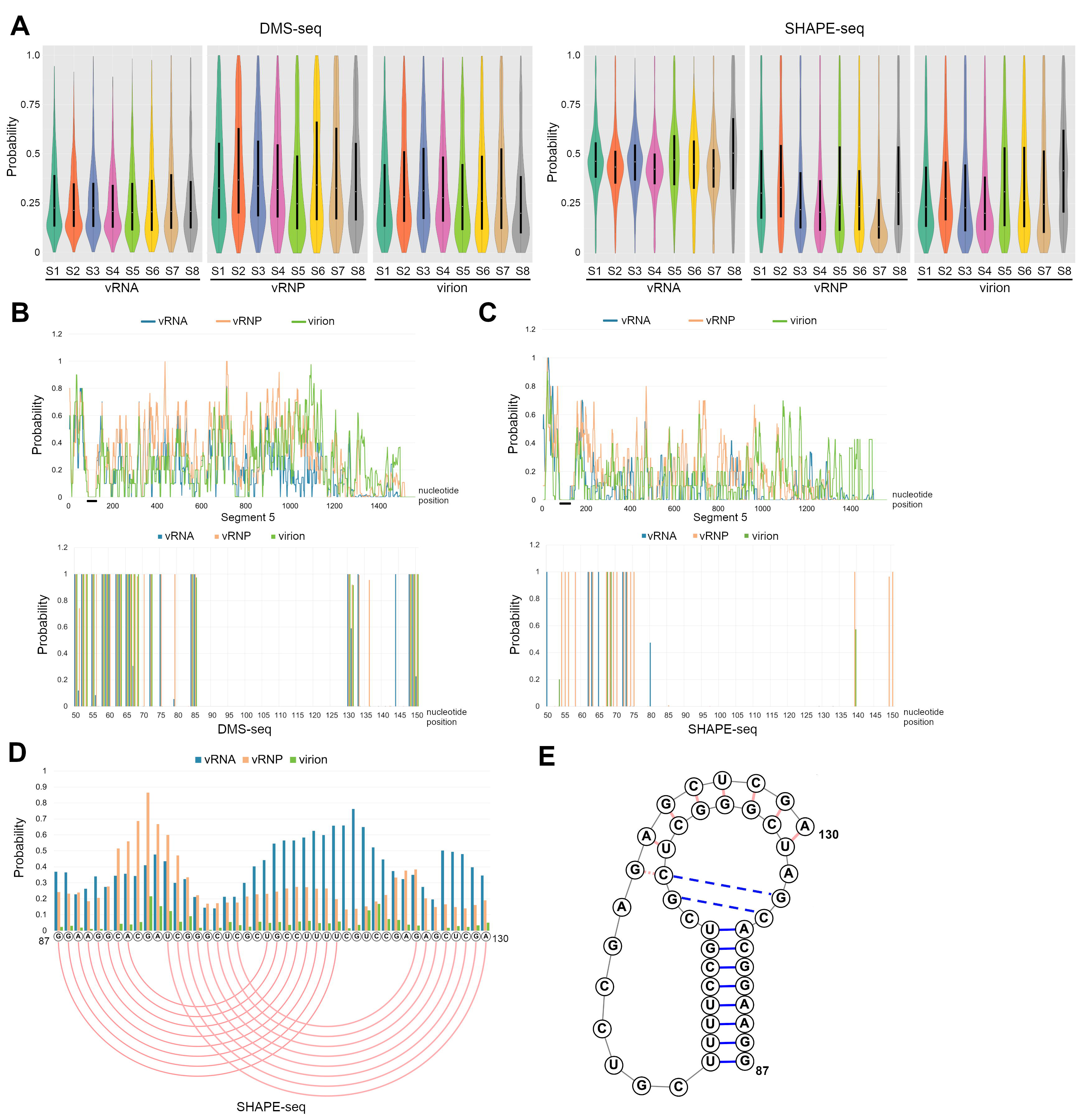
A pseudoknot structure at nucleotide positions 87–130 of segment 5 in the virion. (A) Distribution of probabilities from DMS-seq and SHAPE-seq of the vRNA, vRNP, and purified virion from the allantoic fluid. Probabilities from DMS-seq and SHAPE-seq were calculated by reactIDR, and the scores of each segment were shown by violin plot. (B and C) The probabilities of segment 5 in the vRNA, vRNP, and virion. Probabilities were calculated by BUMHMM, and a 10-nt moving average of the probability of segment 5 from DMS-seq (B) and SHAPE-seq (C) was shown (upper panels). The lower panels show the probabilities of a specific region indicated by a black line in the upper panels. vRNA sequence is numbered from 5′ to 3′. (D) The probabilities and predicted base pairs at nucleotide positions 87 – 130 of segment 5. The probabilities of each nucleotide from SHAPE-seq were calculated by reactIDR. Pink lines indicate predicted base pairs in the pseudoknot structure. (E) The schematic representation of the pseudoknot structure at nucleotide positions 87 – 130 of segment 5.

We next examined the local RNA secondary structure on vRNP. Base-pairing probabilities from DMS-seq and SHAPE-seq were analyzed by Superfold [40]. Secondary structures on segments 1, 3, 4, and 8 in virion previously identified by SHAPE-MaP analysis [19] were also identified by our DMS-seq and SHAPE-seq (Figure S2A). To identify other structured regions on vRNP, the probabilities were re-calculated by BUMHMM, which outputs the probabilities of modifications of each nucleotide displaying an almost binary output. The probability plot of each segment was shown in Figures 1B, 1C, and S3. We found that the less reactive region for both NAI and DMS was found in vRNA at nucleotide positions 87 – 130 of segment 5 (Figures 1B and 1C). This less reactive region was consistently observed in the vRNP and virion, suggesting that the secondary structure of this region is not changed by the vRNP formation. Interestingly, this region was categorized in the low-NP-binding regions [29] and predicted to form a pseudoknot structure in the previous studies [24, 29]. The base-pairing probability of nucleotide positions 87 – 130 of segment 5 showed that this region could form complex RNA structures (Figure S2B). We tried to determine the RNA structure at nucleotide positions 87 – 130 of segment 5 by RNA structure prediction and SHAPE-seq data. We analyzed the RNA structure of nucleotide positions 87 – 130 of segment 5 only from SHAPE-seq data because DMS labeled only adenine and cytosine residues, and thus the resolution of DMS-seq was lower than that of SHAPE-seq in the region. Figures 1D and 1E show a predicted secondary structure by IPknot [41] for the 87 – 130 nucleotides of segment 5 and the probabilities of the NAI-labeled vRNA, vRNP, and virion at each nucleotide position calculated by reactIDR. The location of predicted pseudoknot structure by IPknot was inconsistent with SHAPE-MaP data [19]. However, our reanalysis from the previous SHAPE-MaP suggests that nucleotide positions 87 – 130 form pseudoknot structure predicted by IPknot (Figure S4). The probabilities of the NAI- labeled vRNP and virion at nucleotide positions 87 – 130 supported the stem and loop structure predicted by IPknot while the probability in the vRNA only weakly supported the stem and loop structure, suggesting that the pseudoknot structure at nucleotide positions 83 – 130 is more frequently formed in the vRNP and virion. Taken together, our comprehensive RNA structural analysis indicated the formation of the particular pseudoknot structure on segment 5 vRNP.

### The RNA structure at nucleotide positions 87 – 130 of segment 5 in mutant viruses

To investigate the role of the pseudoknot structure for virus propagation, we constructed recombinant viruses where multiple mutations were introduced; one is to disrupt the pseudoknot structure (referred as 87mut, hereafter) and another is to reconstruct the base pairs disrupted by the mutations in 87mut (referred as 87rec, hereafter) (Figure 2A). The 87mut virus had three mutations, G96A, C126U, and G129A, that did not induce amino acid changes of NP coded on segment 5, and the 87rec virus had additional three mutations, C98A, G102A, and C105U, that were thought to revert base pairs disrupted by mutations in 87mut virus and that also did not induce amino acid changes of NP (Figure 2A). Among the three mutations within the pseudoknot region, G96 is located within the loop in our predicted pseudoknot structure but within the stem region in a slightly different pseudoknot structure predicted in a previous study [24]. We first analyzed whether these mutations affected the virus propagation. As a result, the propagation of the 87mut virus was impaired compared with that of the wild type virus, even though all three mutations did not change any amino acid residues (Figure 2B). The propagation of the 87rec virus was comparable with that of the wild type virus (Figure 2B). These results suggest that the pseudoknot structure in segment 5 plays an important role in virus propagation.

**Figure 2.**
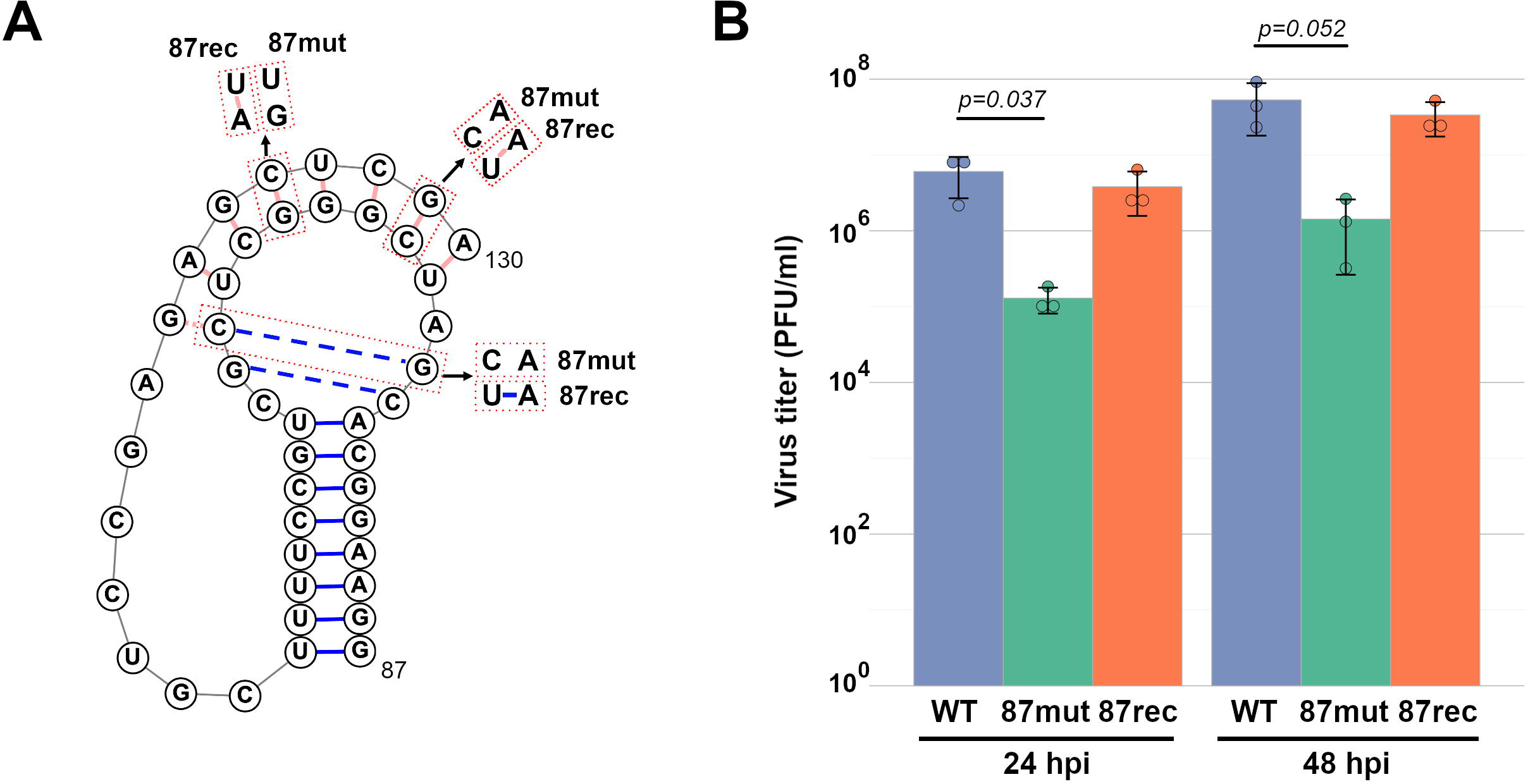
Impairment of propagation of recombinant virus which had mutations in the pseudoknot structure. (A) Mutations in the 87mut and 87rec viruses. The orange boxes indicate the mutated base pairs in the 87mut and 87rec viruses. (B) Virus propagation of the 87mut and 87rec viruses. MDCK cells were infected with the wild type, 87mut, or 87rec virus at an MOI of 0.01. The supernatant was collected at indicated hours post infection (hpi), and the virus titer was determined by a plaque assay. The graph indicates average values with standard deviations from three independent experiments. The circles indicate the titer of each experiment. P-values were calculated by the Dunnett’s multiple comparison test.

Next, we examined the structural differences using SHAPE-seq for the 87mut and 87rec viruses. The wild type, 87mut, and 87rec viruses were purified from infected cell culture supernatant, and SHAPE-seq was performed. The coverages of duplicate experiments were shown in Figure S5A, and the plots of drop-off rate of duplicate experiments and the coefficient of determination were shown in Figure S5B. The coverages were enough to calculate the reactivities except for the 3′ end of segments and the coefficient of determination of each duplicate experiment ranged from 0.40 to 0.94. The probabilities from duplicate experiments were calculated by reactIDR. First, to confirm the formation of the pseudoknot structure in the wild type virus from different virus sources, probabilities at nucleotide positions 87 – 130 of segment 5 in the wild type virus from cell culture supernatant were compared with those from allantoic fluid. The pattern of probabilities in the wild type virus from cell culture supernatant fits the predicted pseudoknot structure, even though the background probabilities were higher than that from allantoic fluid (Figures 1C and 3A). The total probabilities between virus from cell culture supernatant and allantoic fluid were correlated (Figure S6). These results suggest that the secondary structure at nucleotide positions 87 – 130 of segment 5 from different virus sources is almost identical.

**Figure 3.**
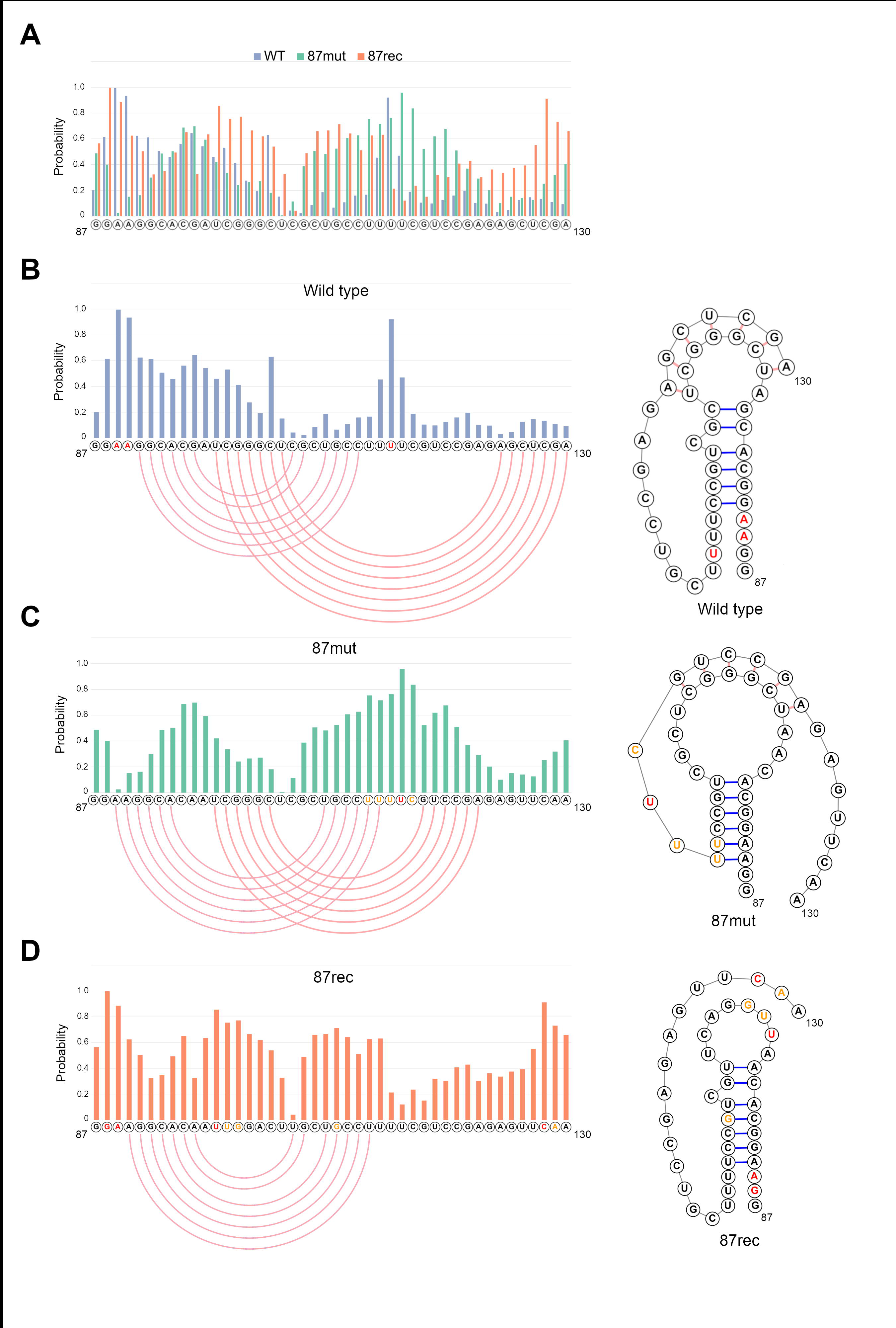
Rearrangement of the RNA structure at nucleotide positions 87 – 130 of segment 5 in the 87mut and 87rec viruses. (A) Probabilities at nucleotide positions 87 – 130 of segment 5 in the wild type, 87mut, and 87rec viruses. Probabilities from SHAPE-seq of the wild type, 87mut, and 87rec viruses were calculated by reactIDR. (B, C, and D) Probabilities and predicted secondary structure at nucleotide positions 87 – 130 of segment 5. Probabilities at nucleotide positions 87 – 130 of segment 5 in the wild type (B), 87mut (C), or 87rec virus (D) are shown, and the secondary structure was predicted by IPknot or MXfold2. Pink lines indicate the predicted base pairs. Red letters and yellow letters indicate probabilities more than 0.85 and 0.70, respectively.

The secondary structure at nucleotide positions 87 – 130 in the 87mut virus was predicted by IPknot. In the 87rec virus, a pseudoknot structure was not predicted at nucleotide positions 87 – 130 by IPknot. Thus, the secondary structure at this region was predicted by MXfold2 [42]. The probability at nucleotide positions 87 – 130 in the wild type, 87mut, and 87rec viruses was shown in Figure 3A. The RNA structure predicted by IPknot (wild type and 87mut viruses) or MXfold2 (87rec virus) and the probabilities of each nucleotide were shown in Figure 3B-D, respectively. In nucleotide positions 87 – 130 of 87mut virus, a pseudoknot structure was also predicted by IPknot. However, the base pairs were substantially reorganized compared with that of the wild type virus. The probability of this region in the 87mut virus corresponded to the predicted pseudoknot structure (Figure 3C). In the 87rec virus, a pseudoknot structure was not predicted in the region by IPknot, while a stem-loop structure was predicted by MXfold2. The pattern of probability at nucleotide positions 87 – 130 of 87rec virus also indicated that base pairs were not formed in the loop region (Figure 3D). These results suggest that the RNA structure of nucleotide positions 87 – 130 is substantially reorganized in 87mut virus and partially reconstituted in 87rec virus.

### Impairment of viral genome replication by mutations in pseudoknot structure in segment 5

To assess in which step at which virus propagation was impaired in the 87mut virus, we determined the amounts of the vRNA segment in the cells infected with the mutant viruses. The relative amount of vRNA in the cells infected with the 87mut virus was decreased at 8 hours post-infection (hpi) and 16 hpi but not statistically significant, while that with the 87rec virus was comparable with that with the wild type virus (Figure 4A). To analyze the replication efficiency of each segment, the ratio of each segment at 8 and 16 hpi was determined by the normalization by the amount of segment 5 (Figure 4B). The relative amount of segment 3 in cells infected with the 87mut virus at 8 hpi was decreased, while that with the 87rec virus was comparable with that with wild type virus. The relative amount of segments in cells infected with mutant viruses at 16 hpi followed the same trend as that at 8 hpi. These results suggest that the mutations in the pseudoknot region affect the replication of vRNAs and the replication ratio of segments, especially segment 3.

**Figure 4.**
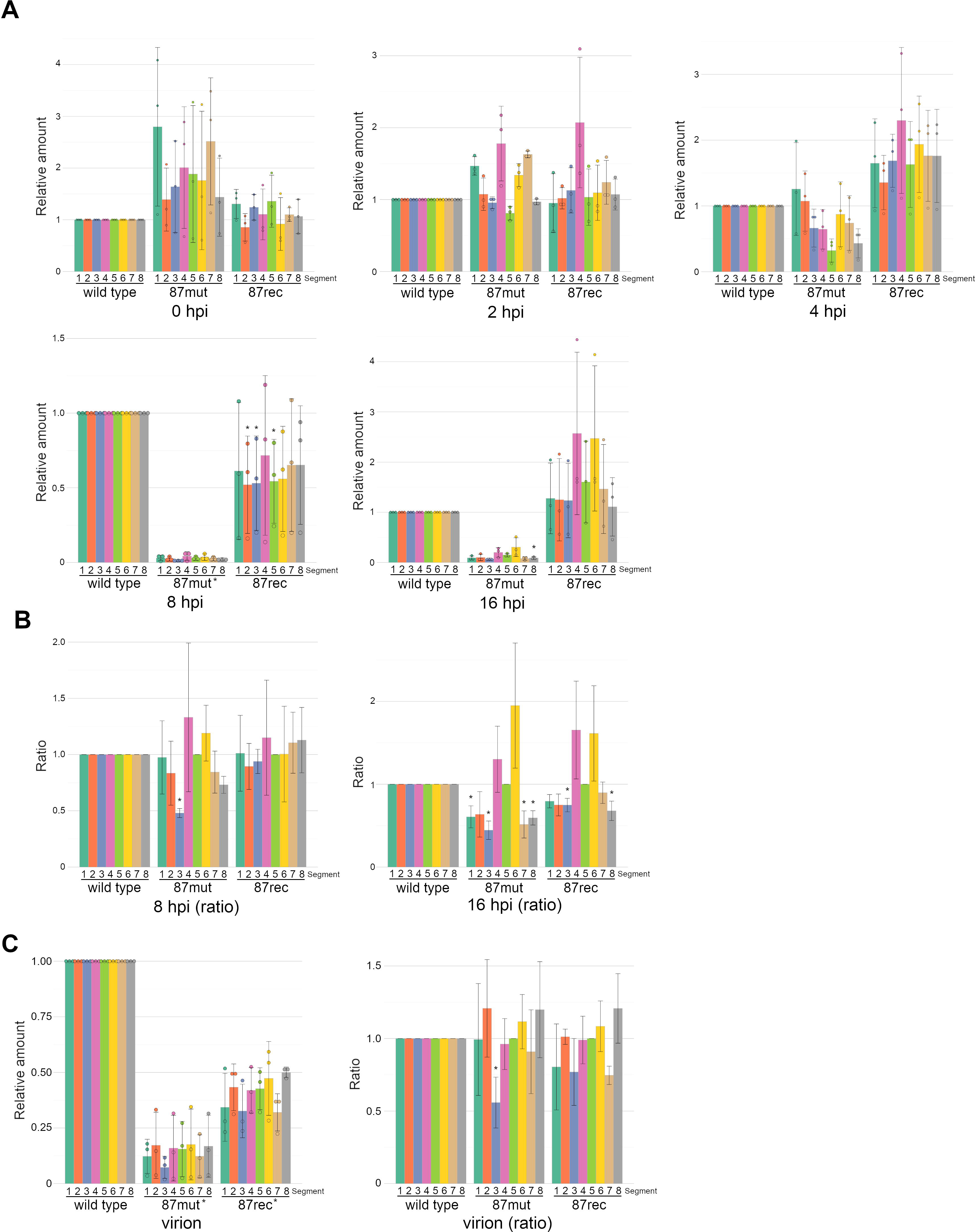
Impairment of viral genome replication and packaging of segment 3 in the 87mut virus. (A) Relative vRNA amount in the infected cells. The amount of each segment was determined by RT-qPCR, and the relative amount was calculated by normalization to the wild type virus. The graph indicates average values with standard deviations from three independent experiments. The circles indicate the relative amount of segments in each experiment. P-values were calculated by the Dunnett’s multiple comparison test, and an asterisk indicates P-values less than 0.05. An asterisk beside 87mut at 8 hpi result means P-values of all segments less than 0.05. (B) The ratio of segments in infected cells. The relative vRNA amount at 8 hpi and 16 hpi in (A) was double-normalized by the amount of segment 5 to calculate the ratio of segments in infected cells. P-values were calculated by the Dunnett’s multiple comparison test, and an asterisk indicates P-values less than 0.05. (C) Relative vRNA amount and ratio of segments in the virion. The amount of each segment was determined by RT-qPCR, and the relative vRNA amount was calculated by normalization to the wild type virus (left graph). The graph indicates average values with standard deviations from three independent experiments. The circles indicate the relative amount of segments in each experiment. The relative vRNA amount was double-normalized by segment 5 to calculate the ratio of segments in virion (right graph). P-values were calculated by the Dunnett’s multiple comparison test, and an asterisk indicates P-values less than 0.05. Asterisks beside 87mut and 87rec mean P-values of all segments less than 0.05.

Furthermore, to analyze the vRNA packaging efficiency of the mutant virus, we determined the amount of vRNAs and the ratio of segments in the virion. Consequently, the amount of vRNAs and the relative amounts of segment 3 in the 87mut virus were significantly decreased than that of the wild type virus (Figure 4C). These results suggest that the packaging of segment 3 was impaired depended on the amount and ratio of segments in infected cells. Moreover, we performed a FACS analysis of the viral proteins in the cells infected with the mutant virus to quantify the fraction of semi-infectious particles that represented the segment bundling function [43]. The infected cells were stained with the combination of NP and M1, HA and NP, and HA and M1, respectively. In cells infected with the mutant viruses, the ratios of the cells expressing NP and M1, HA and NP, and HA and M1 were comparable to that of the cells infected with the wild type virus (Figure S7). This result indicates that the co-packaging efficiency of these segments is not altered in mutant viruses.

### Intersegment structures supported by RNA interactions

We showed that the packaging of segment 3 in the 87mut virus was significantly decreased, and it has been provided evidence that intersegment RNA interactions drive segment bunding. Thus, we next analyzed comprehensive intersegment interactions in the wild type and the mutant viruses. To identify global genome RNA interactions in the virion, we optimized a LIGR-seq, which cross-links RNAs that form base pairs between each other, for the virion [39] (Figure S8). The total number of paired-end reads mapped at intrasegment and at intersegment in each LIGR-seq were listed in Table S3. First, the consistency of intrasegment interaction detection is investigated using our modified LIGR-seq. To quantify the interaction frequencies, the contact map was normalized using the iterative method that was employed in Hi-C data analysis [44]. The normalized count in each 100 nt bin was referred to as the contact score. To assess a bias of selection of the cross-linked RNA, we performed LIGR-seq experiments with or without RNaseR treatment. Background intrasegment signals were reduced in the RNaseR-treated samples, and the intersegment signal was enhanced (Figure S9). To identify the reliable intersegment interactions, the contact scores from the duplicate experiment were adjusted using the irreproducible discovery rate (IDR) [45]. We determined the threshold to the IDR score of the 100th intersegment interaction (Figure S10). Regions with high IDR scores were mainly intrasegment interaction regions, but reproducible intersegment interaction regions were identified by the IDR analysis (Figure S10). We constructed a contact map of intersegment and intrasegment interactions and an interaction map of identified 100 intersegment interactions (Figure 5). As a result, the interaction of the 3′- and 5′- end of the vRNA was captured (Figure 5A), and intersegment interactions formed redundant and complex interaction networks (Figure 5B).

**Figure 5.**
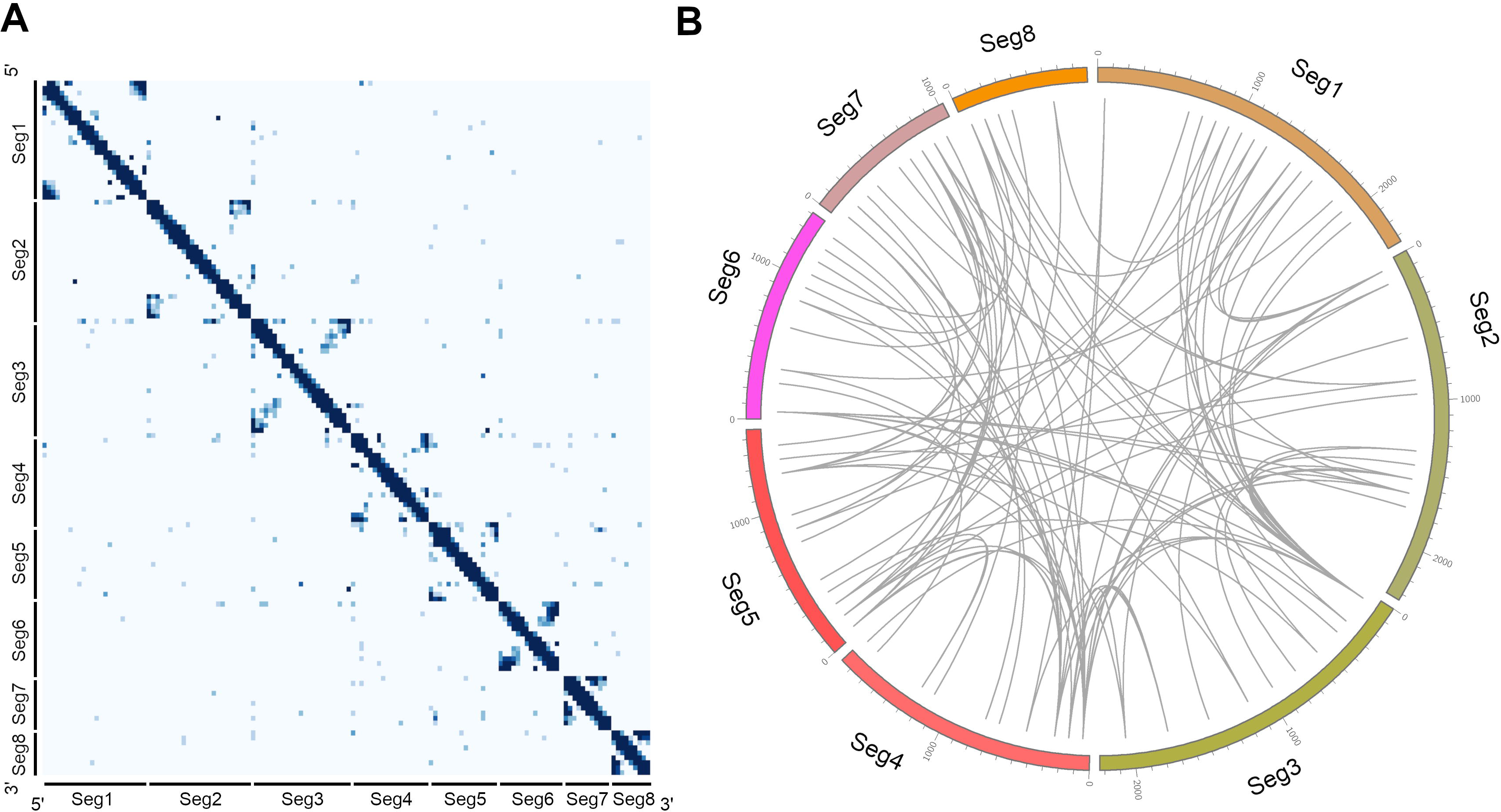
Intra- and intersegment interaction map of segment RNAs. **(A) Contact** map of the wild type virus by duplicate LIGR-seq. Intrasegment and intersegment interactions were identified by IDR score (IDR score > 0.01, containing 100 intersegment interactions) from duplicate LIGR-seq data, and contact maps were constructed. The light and shade of colors in each bin represent low and high normalized contact scores. Bin: 100 bp. (B) The intersegment interaction map in the purified virion. Intersegment interactions with IDR scores from the top to the 100th were extracted. Each line indicates the intersegment interaction which results from LIGR-seq.

Moreover, to confirm the reproducibility of the intersegment interactions captured by LIGR-seq, we compared our global map of intersegment interactions with those previously identified by sequencing of psoralen-crosslinked, ligated, and selected hybrids (SPLASH) [19]. The global map of intersegment interactions was reported in PR8 and WSN strain by SPLASH [19] and WSN strain by dual cross-linking, immunoprecipitation, and proximity ligation [18]. We carried out LIGR-seq analysis on the PR8 strain, and intersegment interaction analysis by SPLASH suggested that the prevalence of the intersegment interactions was prone to change between PR8 and WSN strain [19]. Thus, we compared the 100 intersegment interactions of LIGR-seq and the intersegment interactions with the top to the 100th read score of SPLASH analysis. Intersegment interaction maps from our modified LIGR-seq and SPLASH are shown in Figure S11. We defined the intersegment interactions within 200 nt of each other as overlapped intersegment interactions because vRNA was digested into a fragment that was approximately 300 nt long in our library preparation for large-scale sequencing. Twenty-six intersegment interactions (26% of the 100 interactions detected by LIGR-seq) were identified in both LIGR-seq and SPLASH, suggesting that intersegment interactions captured by LIGR-seq are also captured by a different method, partially.

### Rearrangement of the intersegment interaction of segment 3 in the 87mut virus

To analyze whether intersegment interactions were rearranged in the mutant viruses, we further performed LIGR-seq for the 87mut and 87rec viruses and obtained the intersegment interaction map. First, we analyzed the intersegment interactions identified in both the wild type and the mutant viruses. As a result, 20 interactions in the 87mut virus and 37 interactions in the 87rec virus were overlapped with the interactions in the wild type virus (Figure 6A, red line). The interactions between segments 3, 4, 6, and 8 and other segments identified both in the wild type and the 87mut virus were few, while a novel interaction was observed in the 87mut virus (Figures 6A and S12A). The bias to specific segments was not observed in the interactions identified both in the wild type and the 87rec virus (Figure 6A and S12B). At the nucleotide positions 1 – 200 of segment 5 where mutations were introduced in the mutant viruses, the interactions between segments 5 and 4 were not maintained in the 87mut virus (Figure S12C). The interactions between nucleotide positions 1 – 100 of segment 5 and segments 4 and 7 were maintained in the 87rec virus, while those between segment 5 and segments 1 and 2 were not maintained (Figure S12C). These results suggest that the intersegment interactions were partially rearranged in the 87mut virus and were maintained in the 87rec virus compared with that in the 87mut virus.

**Figure 6.**
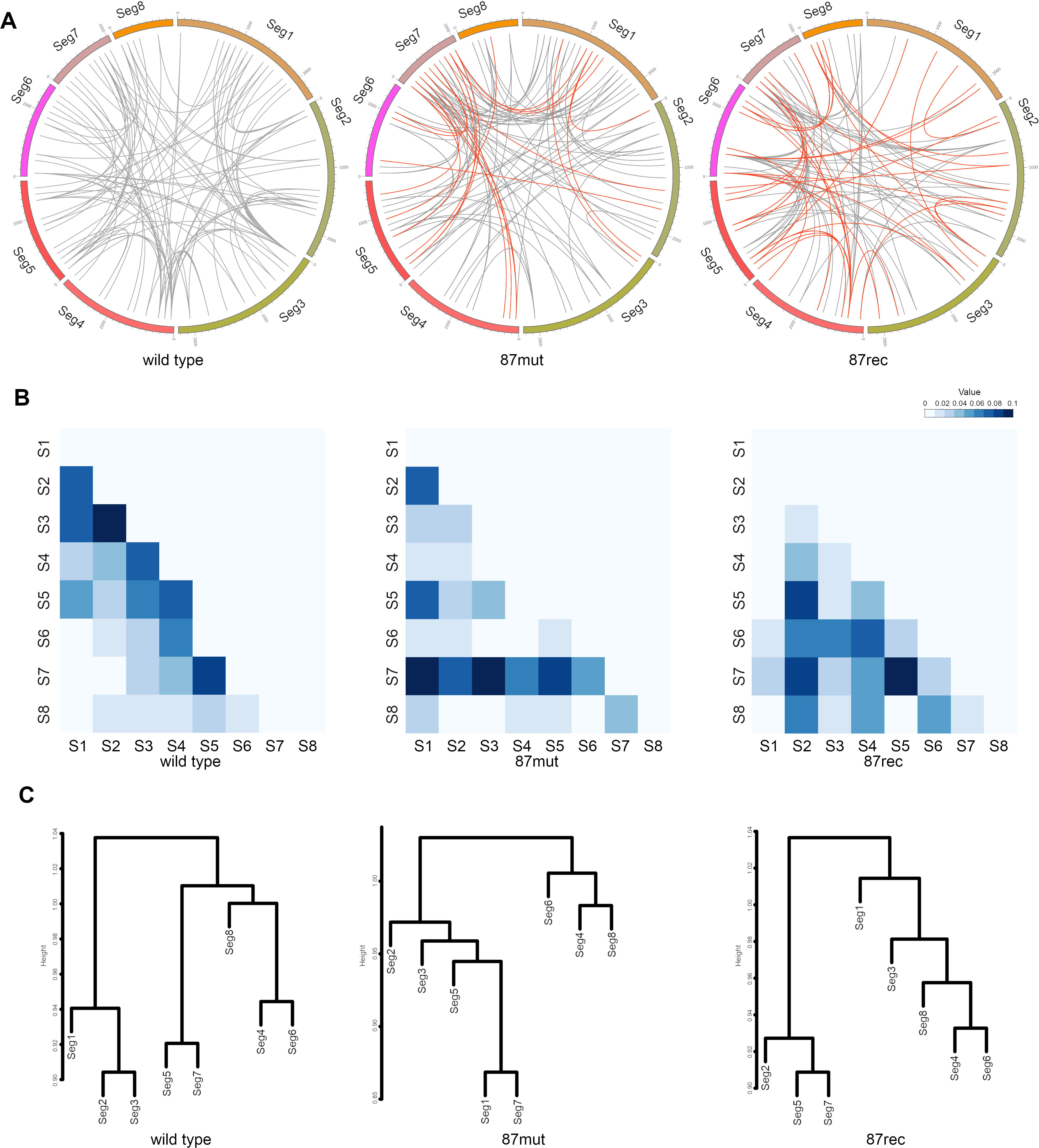
Reconstitution of the intersegment interactions in the 87mut virus. **(A)** Intersegment interactions in the wild type, 87mut, and 87rec viruses. Intersegment interaction maps of the wild type, 87mut, and 87rec viruses were constructed from LIGR-seq. Intersegment interactions identified in both the wild type and the 87mut virus or the 87rec virus within a limit of 200 nt have been indicated by red lines. (B) Intensity maps of the intersegment interactions in the wild type and the mutant viruses. Total contact scores of all the two-segment combinations from LIGR-seq were calculated, and the contact scores were normalized by total contact scores of all intersegment interactions. The light and shade of colors in each bin represent low and high normalized contact scores from 0 to 0.1. (C) Cluster analysis of intersegment interactions from LIGR-seq. Cluster analysis was performed using the reciprocal number of contact scores of all the two-segment combinations from LIGR-seq.

Next, we assessed the intensity of the intersegment interactions in the wild type and the mutant viruses. To analyze the intensity of intersegment interactions, the ratio of the contact scores for all the interactions in each intersegment combination to the total contact scores in all the intersegment interactions was summarized in Figure 6B. First, we focus on the intersegment interactions between segment 3 and other segments because the replication of segment 3 was reduced in cells infected with the 87mut virus. The overall contact scores between segments 3 and other segments except for segment 7 were reduced in the 87mut virus, while that of segment 7 was enhanced. The overall contact scores between segment 3 and segments 6, 7, and 8 were maintained, while those between segment 3 and segments 1, 2, 4, 5 were substantially reduced in the 87rec virus. These results suggest that the intersegment interactions of segment 3 and other segments are rearranged in the mutant viruses. To characterize the specific intersegment interactions in the wild type and the mutant viruses, a cluster analysis of the overall contact scores in each intersegment combination was performed (Figure 6C). In the wild type virus, the clustering analysis found three clusters where the contact scores show similar tendency: segments 1, 2, and 3, segments 5 and 7, and segments 4, 6, and 8. The interactions of any segment of the first cluster and other cluster segments were rearranged in both the 87mut and 87rec viruses, while the connection of the segment pair between the second cluster and the third cluster was maintained. Taken together, our analysis showed the rearrangement of intersegment interactions that occurred mainly in segments 1, 2, and 3 by the conformational changes of the secondary structure in segment 5.

## Discussion

We utilized both DMS-seq and SHAPE-seq to uncover the secondary structures of vRNA with the viral proteins in the virion because of the difference of their individual advantages. The median of probabilities from the virion and the vRNP labeled with DMS was higher than that from the DMS-labeled vRNA (Figure 1A). On the other hand, the median of probabilities labeled with NAI was lower than that from the NAI-labeled vRNA in SHAPE-seq (Figure 1A). The specificity of DMS and NAI to modify moieties of a single-stranded RNA is different. NAI has been shown to modify the 2′-OH group in the ribose backbone, whereas the vRNA has been shown to bind to NP via the phosphodiester backbone [46]. The reactivity of NAI and DMS is affected by not only RNA structure but RNA binding proteins such as NP. The opposite reactivity results of DMS-seq and SHAPE-seq can be explained by the accessibility of DMS and NAI to nucleotides in the RNA-protein complex. The probability of high-NP binding regions in the vRNP and virion was higher than that of the other regions, while the opposite was observed in the vRNA (Table S2). Thus, we conclude that the secondary structure of vRNA is likely to be dissolved by binding NP.

A previous *in silico* analysis showed that nucleotide positions 87 – 130 of segment 5 could form a pseudoknot structure [24] while CLIP analyses showed that this region was a low NP binding region [28, 29]. In addition, SHAPE-MaP analysis showed that this region formed a pseudoknot structure in the virion [19]. We showed a more precise structure of this region in vRNP form by using two comprehensive RNA structural sequencing with robust statistical analysis. While other additional pseudoknot structure regions were predicted in nucleotide positions 397 – 518 of segments 1 and nucleotide positions 804 – 867 of segment 8 [29], we did not detect these regions as pseudoknot structures from our DMS-seq or SHAPE-seq results. Moreover, the RNA secondary structure prediction by IPknot showed that these regions do not form the pseudoknot structure. Thus, these regions might be unlikely to form pseudoknot structures in the virion. Our structural analyses showed that RNA structures could be formed on vRNP but could not identify the precise RNA structure except for the pseudoknot structure on segment 5. One of the reasons why identifying precise vRNA structures in virion is difficult may be diversity in the structure of vRNAs in the virion. In addition, the selected regions within vRNP can interact with multiple regions in other segments (Figure 5A) [18]. These multiple intersegment interactions at the same region also could be explained by the structural diversity of vRNP in the virion. The structural multi-integrated omics analysis in a single virion would be necessary to reveal the global secondary structure of the viral RNA genome.

The secondary structure of the pseudoknot region in segment 5 was considerably different between the wild type and the 87mut viruses (Figure 3), and replication of vRNAs, particularly segment 3 which did not have mutations, was reduced in the cells infected with the 87mut virus (Figure 4A). Reduction of segment 3 replication could induce the reduction of PA mRNA, which is synthesized from segment 3 vRNA and encodes one of the viral polymerase subunits, resulting in the reduction of replication of all segments. Suboptimal codon pairs of viral mRNA reduce mRNA stability and translation efficiency of the deoptimized gene and that IAV with maximized frequencies of CpG dinucleotides in segment 5 showed attenuation in cell culture [47]. CpGs in segment 5 of the 87mut and 87rec viruses are reduced compared to that of the wild type virus, and the average codon pair scores [48] of the 87mut and 87rec viruses are comparable to that of the wild type virus (wild type: 0.0066, 87mut: 0.0081, and 87rec: 0.0076). Thus, the secondary structure changes by introducing synonymous mutations rather than suboptimal codon pairs and the frequency of CpG dinucleotides affects the replication defect. The secondary structure changes could induce NP repositioning. It is possible that repositioning of NP occurs in segment 3 by the effects of reconstitution of the intersegment interactions in the 87mut virus. Further analysis will be required to clarify the detailed molecular mechanism underlying the replication defect of segment 3 in the 87mut virus. The propagation of the 87rec virus was comparable with that of the wild type virus (Figure 2B) though the pseudoknot structure at nucleotide positions 87 – 130 of segment 5 was not formed (Figure 3D). Replication of segments in cells infected with the 87rec virus was delayed, and the amount of vRNAs in the 87rec virus was reduced (Figure 4). These results indicate that the phenotype is not fully complemented in the 87rec virus. The stem-loop structure at nucleotide positions 87 – 115 in the 87rec virus could partially complement the reduction of replication of vRNAs in cells infected with the 87mut virus.

To identify specific intersegment interactions quantitatively, we used the iterative method which is a method for matrix balancing and IDR in our analysis. The contact score of intersegment interactions in our analysis is expected to be more reliable because the contact scores were normalized by the iterative method. The information on the reproducibility was included by IDR to filter out false positives in the final normalized contact scores. We identified a redundant and complex intersegment network (Figure 5). A recent study also revealed redundant and complex networks of RNA-RNA interactions in the IAV by using other global RNA-RNA interaction detection methods [18, 19]. Redundancy of the intersegment interaction could provide the plasticity that tolerates the loss of some intersegment interactions [49]. This plasticity may allow the IAV to escape an established immunity by mutations, while excess interactions may prevent the segment reassortment which also generates the diversity of IAV. Thus, the balance of intersegment interactions may be a factor for determining the diversity of IAV. Our LIGR-seq analysis identified clusters of segment interactions, and these clusters were largely maintained in the 87mut and 87rec viruses (Figure 6C). It has been reported that specific segments play more important roles than the other segments for viral genome packaging [9, 11], and sequential vRNP associations during cytoplasmic transport of viral genome were observed [50, 51]. Our findings raise the possibility that the cluster of segment interactions generates the hierarchy of segments for viral genome packaging.

Nucleotide positions 1 – 200 of segment 5 are one of the hotspots of intersegment interactions, and nucleotide changes in this region decreased the intersegment interactions not only at this site but between other segments (Figure S12C). The finding that nucleotide changes in a hotspot induced a genome-wide rearrangement of intersegment interactions has been reported [18]. Our results indicate that the unwinding of the pseudoknot structure induces a rearrangement of intersegment interactions. The total contact scores of segment 3 in the 87mut virus were decreased, and that of segment 7 was increased (Figure 6B). The intersegment interactions of segment 3 with other segments were equally decreased except for those between segments 3 and 7 (Figure 6C). One possible explanation of this finding is that segment 3 is eliminated from the center of the ‘7+1’ vRNP arrangement in the 87mut virus by disrupting the pseudoknot structure of segment 5. The total contact scores of segments 1 and 3 in the 87rec virus were decreased through the virus propagation of the 87rec virus was comparable with that of the wild type virus (Figure 6B). Nucleotide positions 87 – 130 of segment 5 did not form a pseudoknot structure in the 87rec virus, but nucleotide positions 87 – 115 form a stem-loop structure (Figure 3). This stem-loop structure partially complements not only the reduction of replication of vRNAs but the intersegment interaction rearrangement with a loss in replicative fitness.

Overall, our study presents the global secondary structure and intersegment interactions of the IAV genome in the virion. We showed a functional pseudoknot structure on the vRNP. These findings will help us to understand the molecular mechanisms underlying the emergence of potential pandemic IAV that is generated by segment reassortment will contribute to developing a new class of anti-influenza drugs that bind and unwind the specific RNA structure in the IAV genome.

## Materials and methods

### Cells

MDCK cells were maintained in a minimal essential medium (MEM) (Sigma-Aldrich, ST. Louis, MO) containing 10% fetal bovine serum and penicillin/streptomycin (Nacalai Tesque, Kyoto, Japan). HEK293T cells were maintained in a Dulbecco’s modified Eagle’s medium (DMEM) with high glucose concentration (Sigma-Aldrich) containing 10% fetal bovine serum and penicillin/streptomycin.

### Viruses

Influenza virus A/PR/8/34 (H1N1) (PR8) was grown in the allantoic sacs of 11 days-old chick embryos at 35.5°C for 48 h. The purified virion and the vRNP from the purified virion were prepared as previously described [52]. To construct the pPolI-PR8 mutant vector, an inverted PCR was performed using the pPolI-PR8 segment 5 vector as a template with specific primer sets (Primers used in this study were listed in Table S4). After DpnI treatment, phosphorylation, ligation, and transformation into an *Escherichia coli* Mach1 (Thermo Fisher Scientific) were performed. Recombinant viruses were generated using a reverse genetics approach [53]. Viral protein expression vectors [53] and the viral RNA expression vectors derived from the PR8 strain [54] were transfected to 293T cells. To propagate the recombinant virus, MDCK cells were infected with the recombinant virus at a multiplicity of infection (MOI) of 0.1. At 48 h post infection (hpi), the supernatants were collected, and cell debris were removed by low-speed centrifugation (3k × *g*, 5 min). The virus titer was determined by a plaque assay. To prepare purified virion, MDCK cells were infected with the recombinant virus at an MOI of 0.1, and the supernatant was collected at 48 hpi. After removal of cell debris by low-speed centrifugation (500 × *g*, 5 min) and filtration through a 0.45-µm filter, the supernatant was ultracentrifuged at 100k × *g* for 1.5 h using an SW28 rotor (Beckman Coulter, Brea, CA) at 4°C. The pellet was suspended in PBS(-) and centrifuged on 30% to 60% sucrose gradients in PBS(-) at 100k × *g* for 1.5 h in an SW28 rotor at 4°C. Viral bands were pooled and re-precipitated by centrifugation in PBS(-) at 120k × *g* for 1.5 h in an SW55 rotor (Beckman Coulter) at 4°C. The precipitated virion was suspended in a DMS buffer (40 mM Hepes-NaOH [pH 7.4], 100 mM NaCl, and 0.5 mM MgCl2) or PBS(-) and stored at -80°C until use.

### DMS-seq and SHAPE-seq

The vRNA was prepared by proteinase K treatment of virion purified from the allantoic fluid at 37°C for 30 min in SDS buffer (0.25% SDS and 100 µg/ml proteinase K in PBS(-)) followed by phenol/chloroform extraction. NAI was synthesized using a previously described method [34]. One µl of DMS (Wako Pure Chemical Industries, Osaka, Japan) or 5 µl of NAI was added to the purified virion (5 µl of the purified virion from allantoic fluid or from 80 ml of cell culture supernatant of infected cells), vRNP (25 µl of vRNP fraction), or vRNA (from 5 µl of purified virion) in 100 µl of DMS buffer. After incubation for 5 min (DMS) or 15 min (NAI) at 25°C, 10 µl of 1 M DTT was added to stop the reaction. Then, the RNA was extracted with phenol/chloroform. Sequencing libraries were prepared using a previously described method [55]. Briefly, cDNA was synthesized with random hexamers containing the Illumina adapters at their 5′- ends using ReverTra Ace (Toyobo, Osaka, Japan). The ssDNA linker containing a 5′ phosphate and 3′ C3 spacer was ligated to the synthesized cDNA using 20 U of the Circligase I (Lucigen, Middleton, WI). The resultant cDNA was amplified by an adapter-based PCR using the KAPA HiFi DNA polymerase (Roche, Basel, Switzerland). Sequencing was performed using a MiSeq (Illumina, San Diego, CA) (2 × 75-bp PE) and NovaSeq6000 (Illumina) (2 × 150-bp PE). The sequence data have been deposited in DDBJ Sequence Read Archive (DRA Accession: DRA009494 and DRA012096). Raw reads were cleaned and trimmed with Trimmomatic v0.36 [56], and the cleaned reads were aligned to the A/PR/8/34 genome using bowtie2 with default parameters. We performed duplicate DMS-seq and SHAPE-seq experiments on two independent samples, and the reactivities of each nucleotide were calculated using reactIDR [37] with --DMS option and with a default setting, respectively, and BUMHMM [36] with a default setting. Base-pairing probabilities were calculated by Superfold [40] from the probabilities of reactIDR. Computational prediction of the RNA secondary structure was performed by MXfold2 [42] and IPknot [41].

To analyze the probability of high-NP binding regions, NP PAR-CLIP data sets (PR8 strain) were downloaded from Sequence Read Archive (SRX3545111) and aligned to the PR8 genome using bowtie2 with default parameters. The coverage of each nucleotide of PAR-CLIP and control RNA experiments was calculated by IGV [57]. We normalized the number of coverages per nucleotide to the total number of coverages to yield a normalized coverage ratio from both PAR-CLIP and control RNA sequencing. vRNA nucleotides with fold-change >2 were identified, and the regions were extracted. Due to the number of reads, we used only one dataset of PAR-CLIP and control RNA-seq.

### AMT cross-linking and RNA ligation experimental method

The purified virion in PBS(-) was treated with or without AMT (final concentration 100 µg/ml) and cross-linked using 365 nm UV for 20 min (1.56 J/cm2). AMT was added again at 10 min after UV irradiation. The viral RNA was extracted with phenol/chloroform and digested with the NEBNext Magnesium RNA Fragmentation Module (New England Biolabs, Ipswich, MA) at 94°C for 2.5 min. The fragmented RNA was treated with 15 U of CIAP (Takara Bio, Otsu, Japan) at 37°C for 30 min, and phosphorylation was carried out by the 10 U of T4 polynucleotide kinase (Toyobo) at 37°C for 1 h.

RNA pull-down procedure was performed using a modification of an RNA antisense purification method previously described for long non-coding RNA and influenza virus mRNA [58, 59]. cDNA of the PR8 strain was synthesized with the Uni12 primer using ReverTra Ace. The fragments of each segment were amplified with a linker sequence containing specific primers using Taq polymerase (Roche). Specific primers with a linker sequence were designed to amplify every 120 bp of the coding region with a 15-bp overlapped region (PCR primers were listed in Table S4). To synthesize the biotinylated cDNA probe, a second PCR procedure was performed with the biotinylated linker primer using Taq polymerase. The biotinylated cDNA probe for each segment was mixed with the Dynabeads MyOne Streptavidin C1 (Thermo Fisher Scientific, Waltham, MA) and incubated on a rotating wheel at 37°C for 1 h in a LiCl hybridization buffer (10 mM Tris-HCl [pH 7.9], 500 mM LiCl, 1 mM EDTA, and 0.1 % NP-40). The beads were washed with the LiCl hybridization buffer and 1x SSPE and were suspended and incubated in 0.15 M NaOH for 10 min. After incubation, the mixture was neutralized with 100 mM Tris-HCl (pH 7.9) and 1.25 M AcOH. The beads were washed with 0.1 M NaOH and were suspended in the LiCl hybridization buffer. The cross-linked RNA was boiled at 85°C for 3 min and was added to the beads. The beads were incubated at 55°C in a Thermomixer (Eppendorf, Hamburg, Germany) at 1,500 rpm for 2 h. The beads were washed with 1x and 0.1x SSPE at 55°C. Proximity ligation was performed with 40 U of the T4 RNA ligase I at 16°C for 16 h in an RNA ligase buffer, and the beads were mixed in a Thermomixer at 1,000 rpm for 15 sec every 15 min. The vRNA was eluted with 5 U of DNase I (Takara Bio) at 37°C for 30 min in a DNase buffer (40 mM Tris-HCl [pH 7.5], 8 mM MgCl2, and 5 mM DTT) and was extracted with phenol/chloroform. The eluted vRNA was treated with 10 U of RNaseR (Lucigen) in an RNaseR buffer (20 mM Tris-HCl [pH 8.0], 100 mM KCl, and 0.1 mM MgCl2) at 37 °C for 30 min and was extracted with phenol/chloroform. The cDNA was synthesized with a random hexamer using the SuperScript III (Thermo Fisher Scientific) and the NEBNext mRNA Second Strand Synthesis Module (New England Biolabs). Sequencing libraries were constructed using the KAPA Hyper Prep Kit (Roche) or NEBNext UltraII DNA Library Prep Kit for Illumina (New England Biolabs). Sequencing was performed using a HiSeq2500 sequencer (Illumina) (2 × 100-bp PE) and a NovaSeq6000 sequencer (2 × 150-bp PE). The sequence data have been deposited in DDBJ Sequence Read Archive (DRA Accession: DRA005778, DRA009492, DRA009493, and DRA012096)

### Data analysis for segment interactions

Raw reads were cleaned and trimmed into the first 25 bases with Trimmomatic v0.36 [56]. The cleaned reads were aligned to the A/PR/8/34 genome using bowtie2 with default parameters [60]. Obtained sequencing reads were classified into two categories: intersegment and intrasegment interaction. We detected the former interaction by the paired-end reads that were mapped at two different segments and the latter ones that were mapped to the same segment at an inverted direction and a long insert length (more than 500 nt). The start positions of the selected pair-reads were counted in every 100 nt, and the contact map was constructed for the intra- and intersegment interactions. To normalize the biases, we adapted the iterative method which has been employed for the Hi-C analysis [44]. The raw contact of each bin in the contact map was divided by the sum of the contacts in the whole row and the sum of the contacts in the whole column. This calculation was repeated until it converges. The normalized count in each 100 nt bin was referred to as the contact score. To discriminate accurately between the true and false signals, we performed LIGR-seq in the duplicate experiments and utilized an irreproducible discovery rate (IDR) [45]. IDR compares a pair of ranked lists by contact scores and assigns IDR scores that reflect its reproducibility. The contact scores of all regions containing both intrasegment and intersegment regions of duplicate experiments were analyzed by IDR, and the IDR score of each region was determined. The intersegment interactions were ranked by IDR value, and intersegment interactions with IDR scores from the top to the 100th were used for further analysis. Total contact scores of all the two-segment combinations from LIGR-seq were calculated. To calculate the reciprocal number of contact scores of all the two-segment combinations, 1 was added to the contact scores of all the combinations. Cluster analysis by the Ward method was performed using the reciprocal number as distance.

### RT-qPCR

Total RNA was extracted from MDCK cells infected at an MOI of 1 using the ISOGEN reagent (Nippon Gene, Tokyo, Japan). For the preparation of the vRNA in the supernatant from the infected cells, MDCK cells were infected with the virus at an MOI of 0.1, and the cells were suspended in MEM containing 0.6 µg/ml TPCK-trypsin (Sigma-Aldrich). At 48 hpi, the supernatant was collected, and cell debris was removed by low-speed centrifugation (500 × *g*, 5 min) and filtration through a 0.45-µm filter (EMD Millipore, Billerica, MA). The pre-cleared supernatant was layered on PBS(-) containing 30% sucrose and centrifuged at 130k × *g* for 1.5 h using an SW55 rotor at 4°C. The pellet was suspended in 100 µl of PBS(-). The vRNA was extracted by phenol/chloroform. For RT-qPCR, the cDNA was synthesized with the Uni12 primer using ReverTra Ace. The synthesized cDNA was mixed with the Thunderbird SYBR qPCR mix (Toyobo) and a specific primer set for each segment. The qPCR reactions were performed using a Thermal Cycler Dice Real-Time System TP800 (Takara Bio), and the relative amounts of each segment were calculated.

### FACS analysis

Rabbit polyclonal antibodies against NP [61] and M1 [62] and mouse monoclonal antibody RA5-22 against HA (BEI Resources, NIAID, NIH, Bethesda, MD) and mAb61A5 against NP [63] were used for FACS analysis. Alexa Fluor 488-conjugated anti-mouse IgG and Alexa Fluor 647-conjugated anti-rabbit IgG were purchased from Thermo Fisher Scientific and BioLegend (San Diego, CA), respectively. MDCK cells were infected with the virus at an MOI of 0.01. At 14 hpi, the infected cells were collected by trypsin and fixed with 4% paraformaldehyde at 25°C for 10 min. The fixed cells were permeabilized with 0.2% NP-40 in PBS(-) at 25°C for 15min. The cells were immersed in 0.2% BSA in PBS(-) at 25°C for 1 h and incubated with primary antibodies at 25°C for 1 h. After washing with PBS(-), the cells were incubated with secondary antibodies at 25°C for 1 h. The cells were suspended in PBS(-) and analyzed by the FACS Lyric Flow Cytometer (BD, Franklin Lakes, NJ).

## Acknowledgements

We thank Dr. Yoshihiro Kawaoka (University of Tokyo) for kindly providing plasmids for the reverse genetics system, Dr. Kyosuke Nagata (University of Tsukuba) for kindly providing polyclonal anti-NP antibody, Dr. Fumitaka Momose (Kitasato University) for kindly providing monoclonal anti-NP antibody and plasmids, Dr. Nobuyuki Kobayashi (Nagasaki University) for kindly providing anti-M1 antibody, and Ms. Yukiko Iwata for technical support of experiments. This work was partly performed in the Cooperative Research Project Program of the Medical Institute of Bioregulation, Kyushu University. This work was supported by JSPS KAKENHI Grant Number 25871077, 15K21607, and 19K07598 to N.T., JSPS KAKENHI Grant Number 221S0002 and 16H06279 (PAGS), Japan Program for Infectious Diseases Research and Infrastructure from AMED Grant Number JP20wm0325008 to N.T., Takeda Science Foundation, GSK Japan Research Grant 2016, and the Waksman Foundation of Japan to N.T..

## Supplemental figure

**Figure S1.**
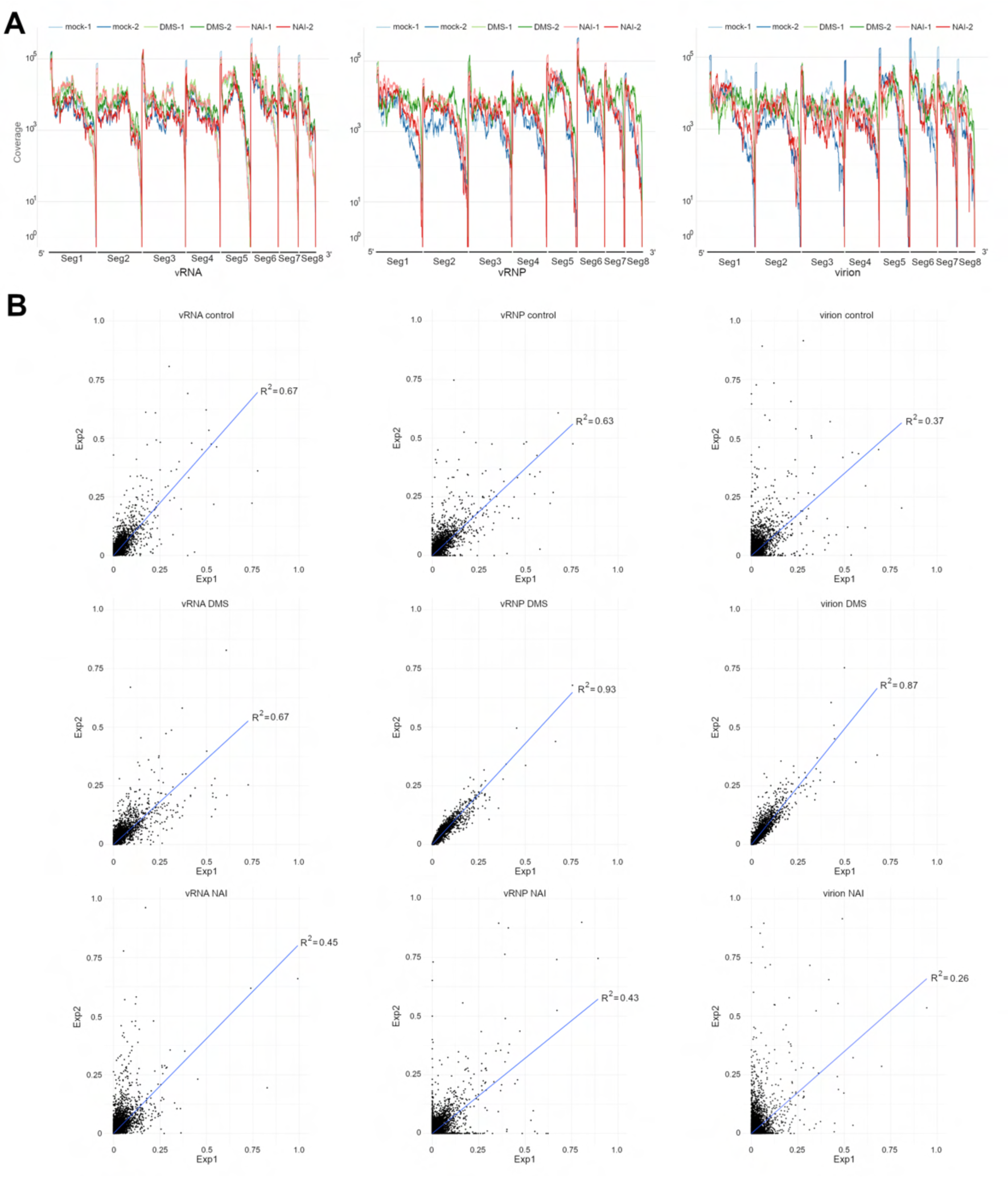
Coverage and drop-off rate of duplicate DMS-seq and SHAPE-seq. (A) Coverage of DMS-seq and SHAPE-seq. Coverages of mock-treated samples, DMS- treated samples, and NAI-treated samples were shown. (B) Scatter plot of drop-off rate from duplicate experiments. The drop-off rate is defined as the value of stopped reverse transcription count divided by the coverage of each nucleotide, and the drop-off rates of duplicate experiments are plotted. The blue line means a regression line derived from the probabilities of each nucleotide. The coefficient of determination (R2) is shown in each graph.

**Figure S2.**
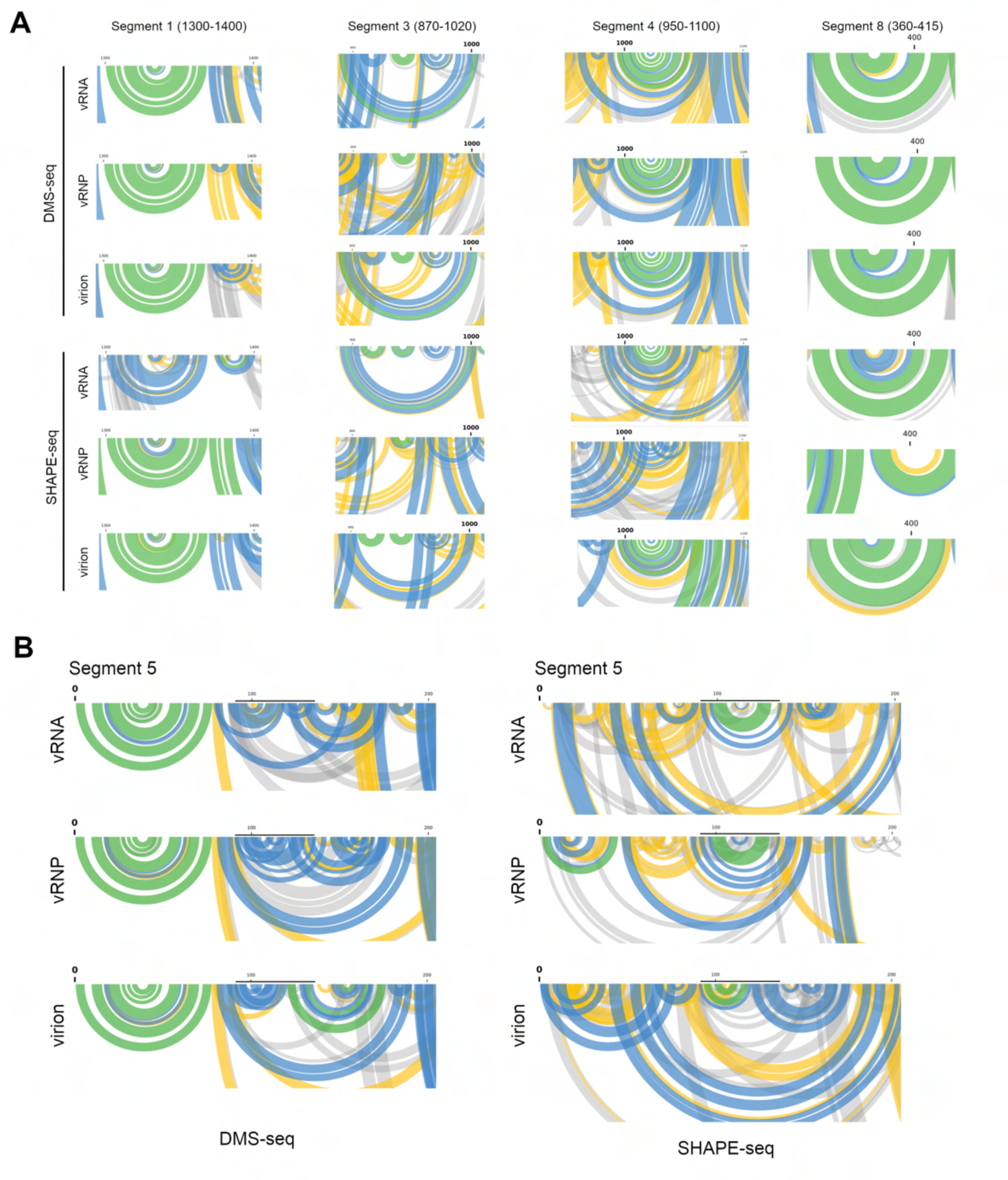
Base-pairing probability from DMS-seq and SHAPE-seq. **Base-pairing** probabilities were calculated from the output of reactIDR by Superfold. Base-pairing probabilities of secondary structure regions identified previous SHAPE-MaP analysis (A) and that of nucleotide positions 1 – 200 of segment 5 (B) were shown. Base pairs were plotted as arc, and green arcs, blue arcs, yellow arcs, and grey arcs indicate a base-pairing probability of 80%, 30%, 10%, 3% or higher, respectively. Bars in (B) indicate nucleotide positions 87 – 130.

**Figure S3.**
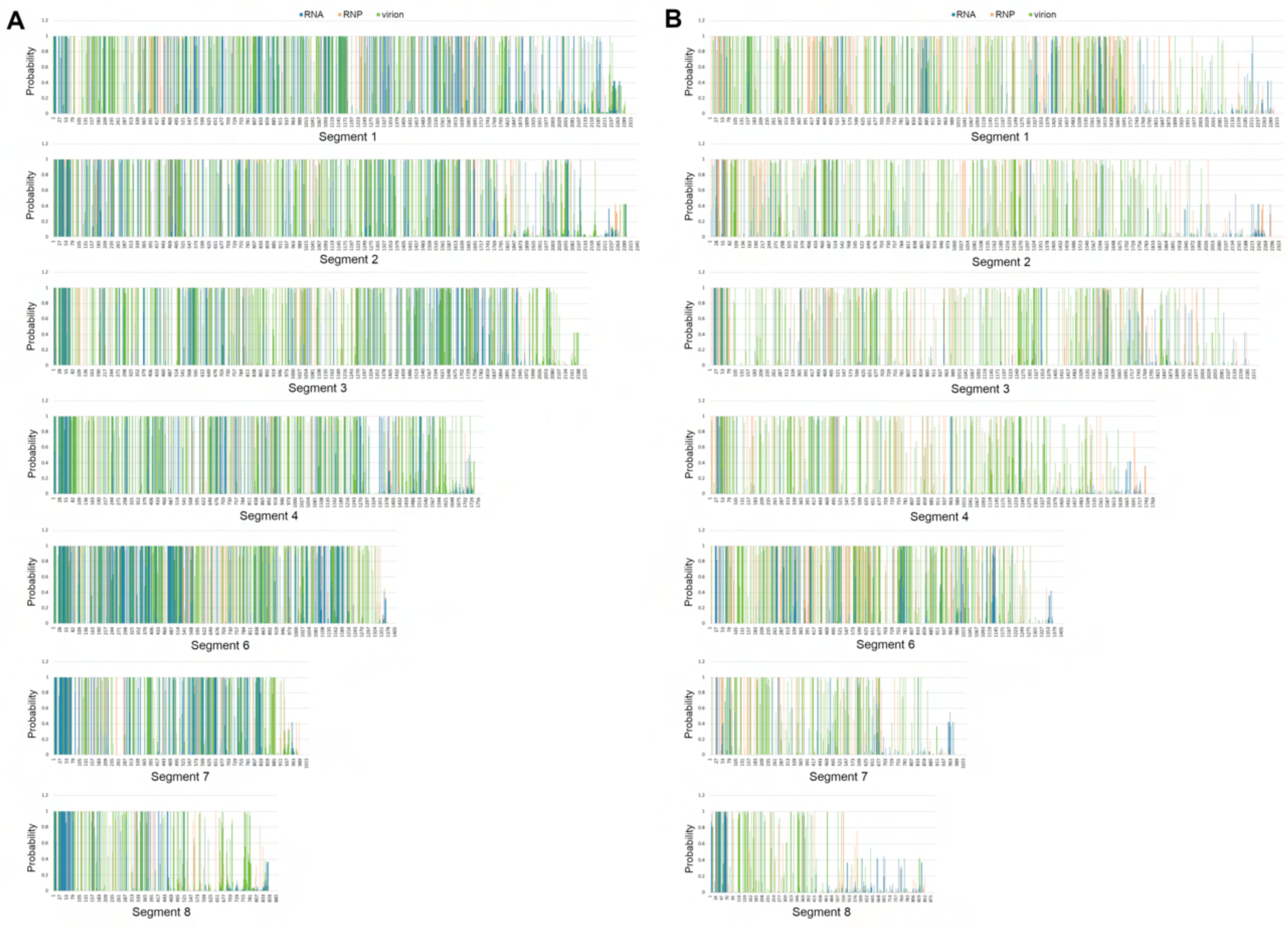
Probabilities from DMS-seq and SHAPE-seq. The probabilities of each segment from DMS-seq (A) and SHAPE-seq (B) were calculated by BUMHMM. Y- axis: probability. X-axis: nucleotide position.

**Figure S4.**
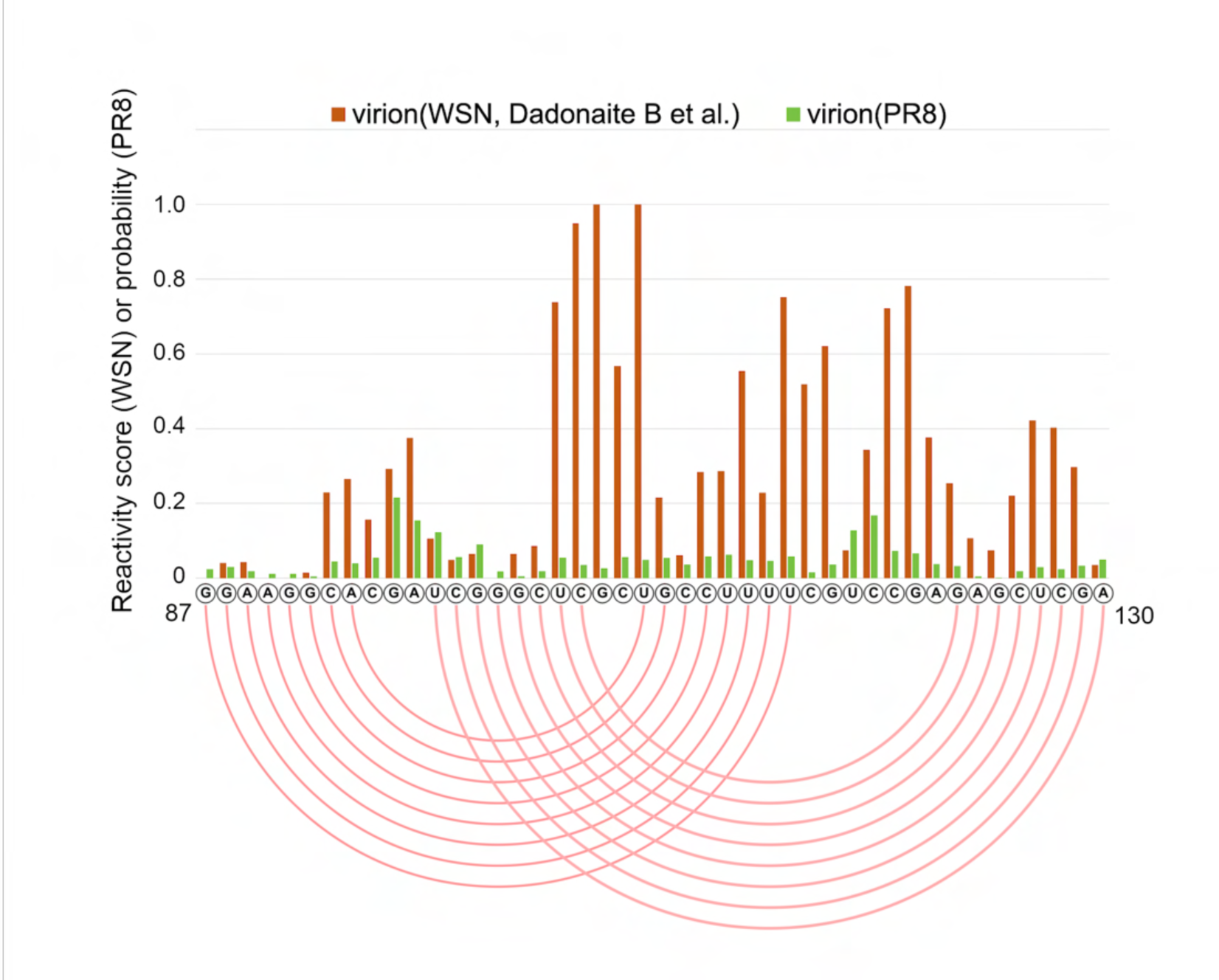
Comparison of the reactivity score at nucleotide positions 87 – 130 of segment 5 by reactIDR with those by the previous study. Reactivity scores at nucleotide positions 87 – 130 of segment 5 from reported SHAPE-MaP (Dadonaite et al., 2019) and probability score of our SHAPE-seq analysis are shown. Pink lines indicate the predicted base pairs from our SHAPE-seq analysis.

**Figure S5.**
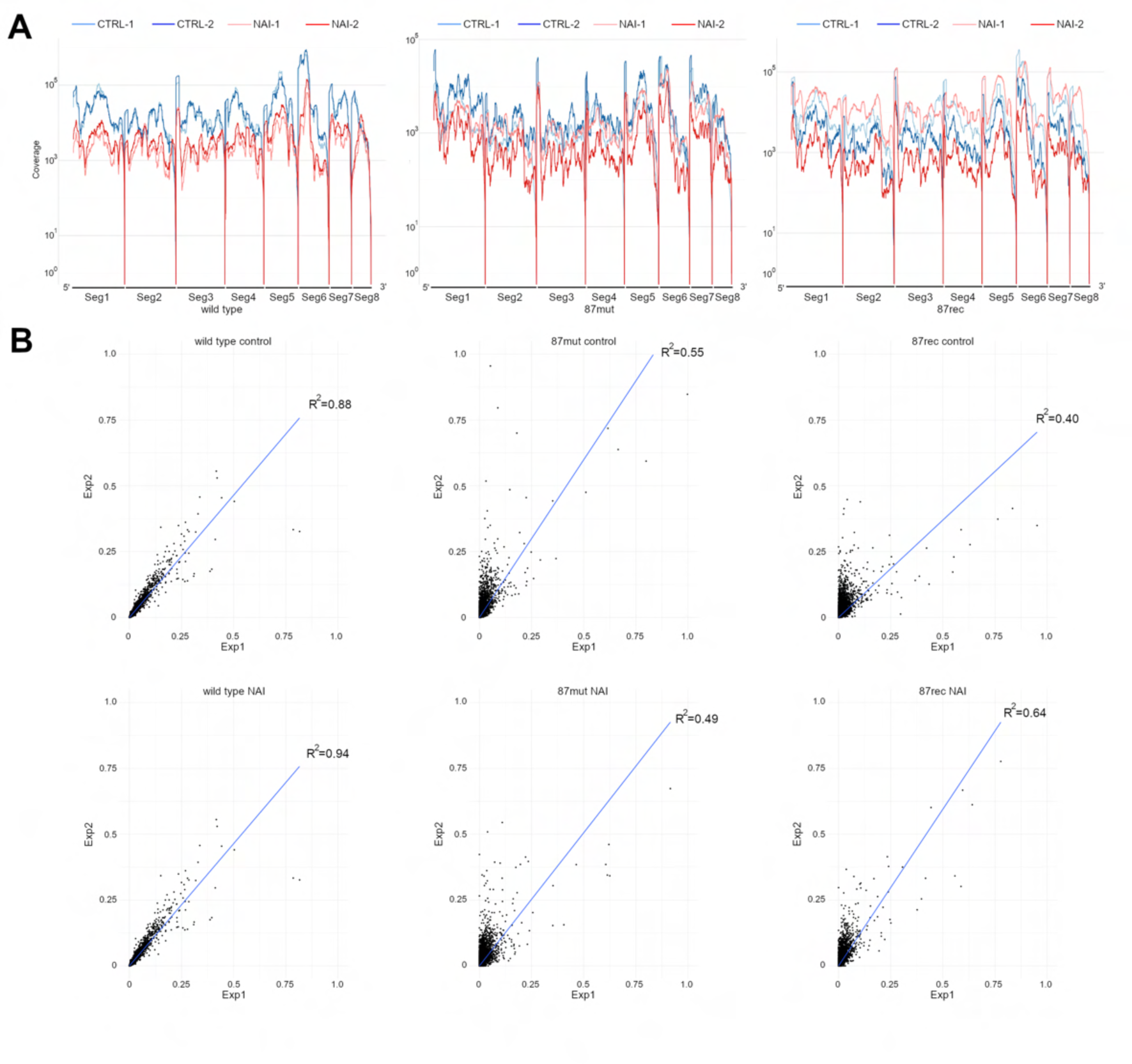
Coverage and drop-off rate of duplicate SHAPE-seq of the wild type, 87mut, and 87rec viruses. (A) Coverage of SHAPE-seq of the wild type, 87mut, and 87rec viruses. Coverages of mock-treated and NAI-treated samples of the wild type, 87mut, and 87rec viruses were shown. (B) Scatter plot of drop-off rate from duplicate experiments. The drop-off rates of duplicate experiments are plotted. The blue line means a regression line derived from the probabilities of each nucleotide. The coefficient of determination (R2) is shown in each graph.

**Figure S6.**
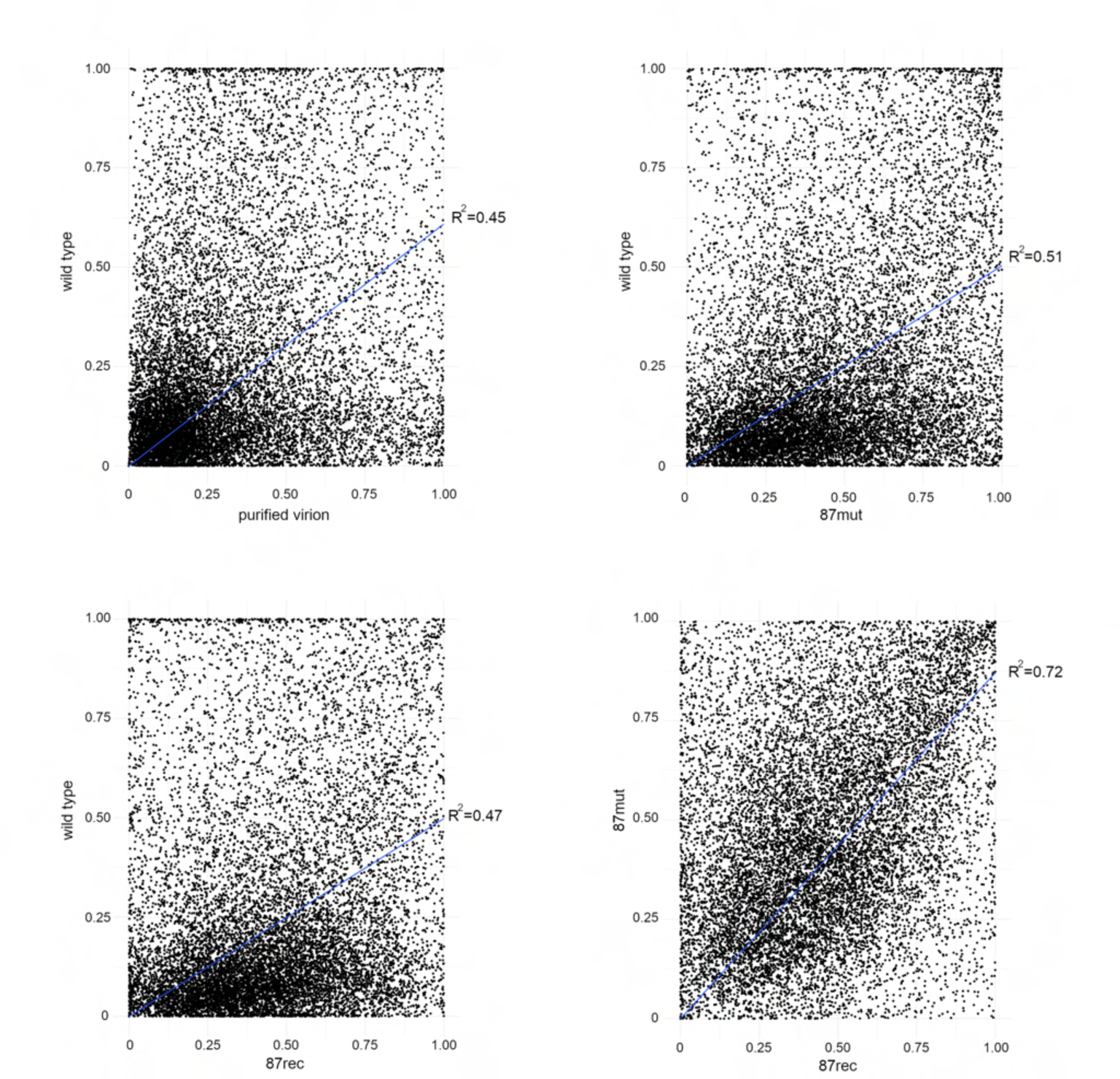
Correlation analyses of probabilities between wild type and mutant viruses. Probability of each nucleotide position calculated from SHAPE-seq of purified wild type virus from cell culture supernatant (wild type) and that from allantoic fluid (allantoic fluid) (upper left), wild type virus and 87mut virus (upper right), wild type virus and 87rec virus (lower left), and 87mut virus and 87rec virus (lower right) are plotted. The blue line means a regression line derived from the probabilities of each nucleotide. The coefficient of determination (R2) is shown in each graph.

**Figure S7.**
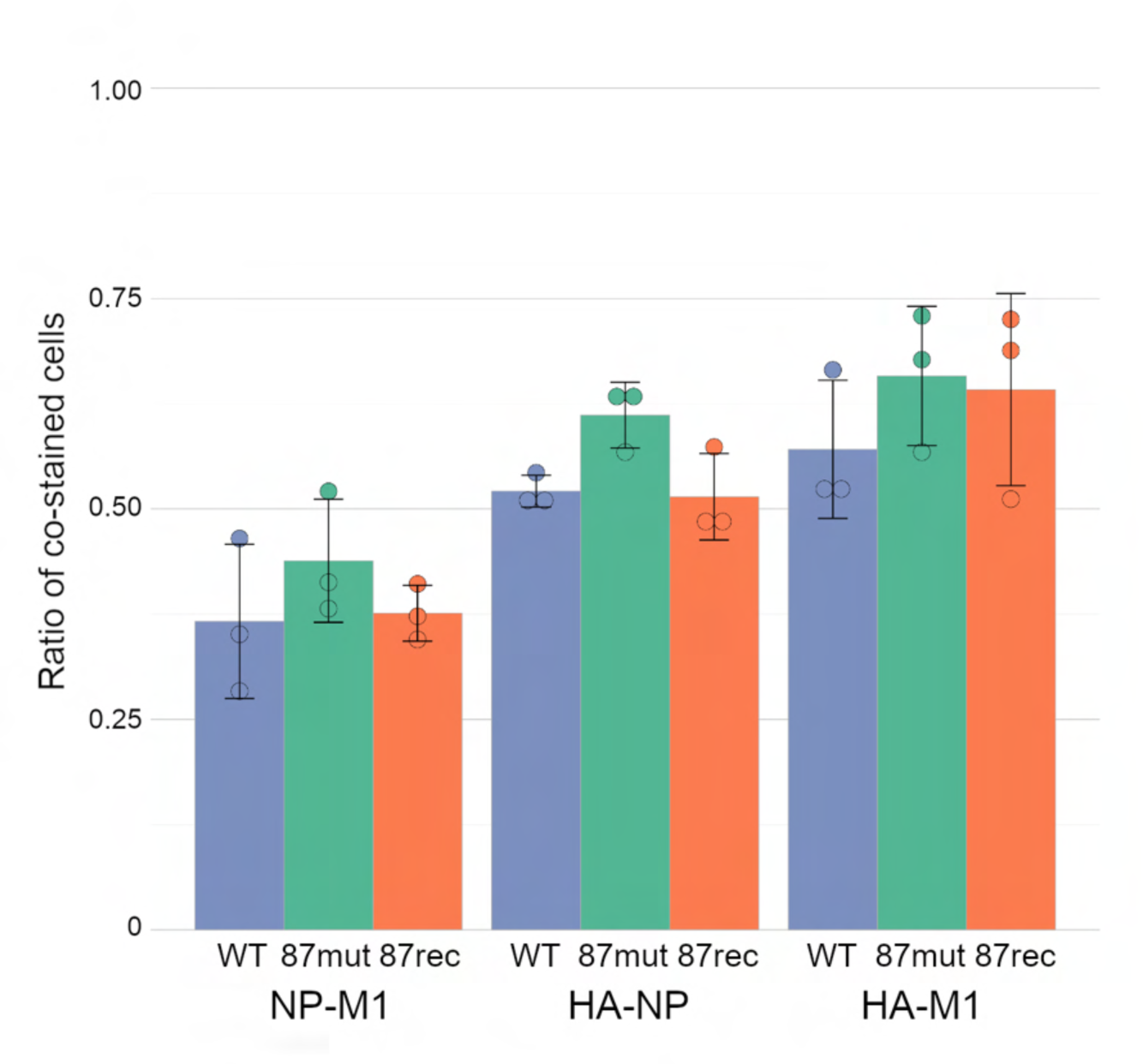
Co-expression rate of two viral proteins in cells infected with the wild type, 87mut, or 87rec virus. The ratio of the co-stained cells was determined by FACS. The graph indicates average values with standard deviations from three independent experiments. The circles indicate the ratio of each experiment.

**Figure S8.**
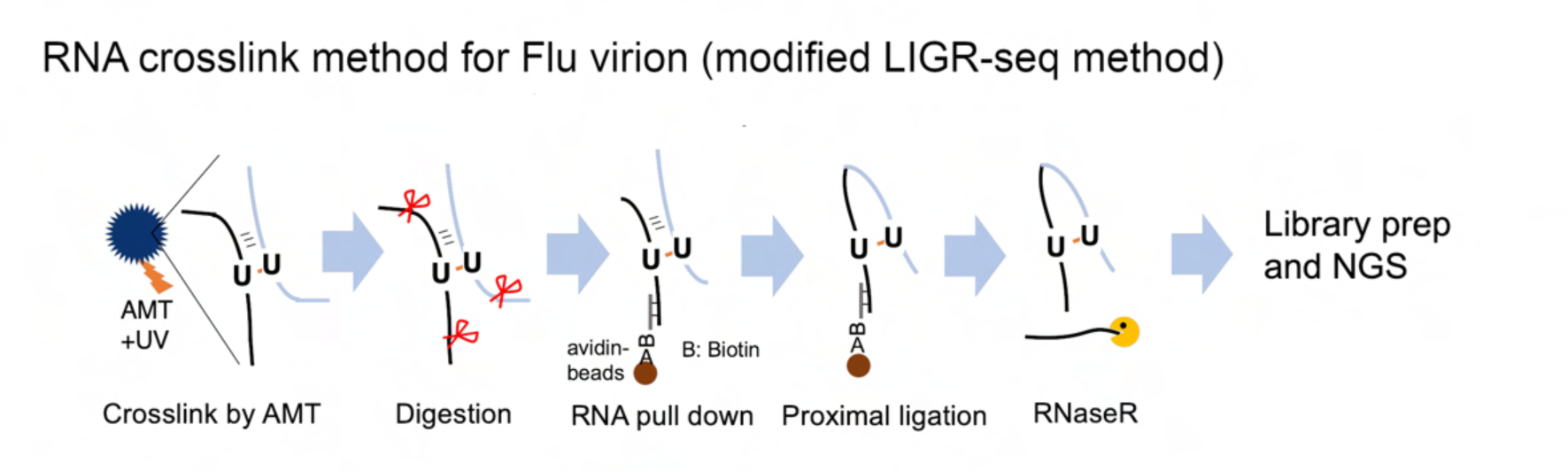
Schematic representation of the methods for the identification of intra- and intersegment interactions by modified LIGR-seq method. Purified virion was treated with AMT to cross-link between RNAs. After partial RNA digestion, vRNA was purified by a biotin-conjugated antisense single-strand DNA, and proximal ligation was performed.

**Figure S9.**
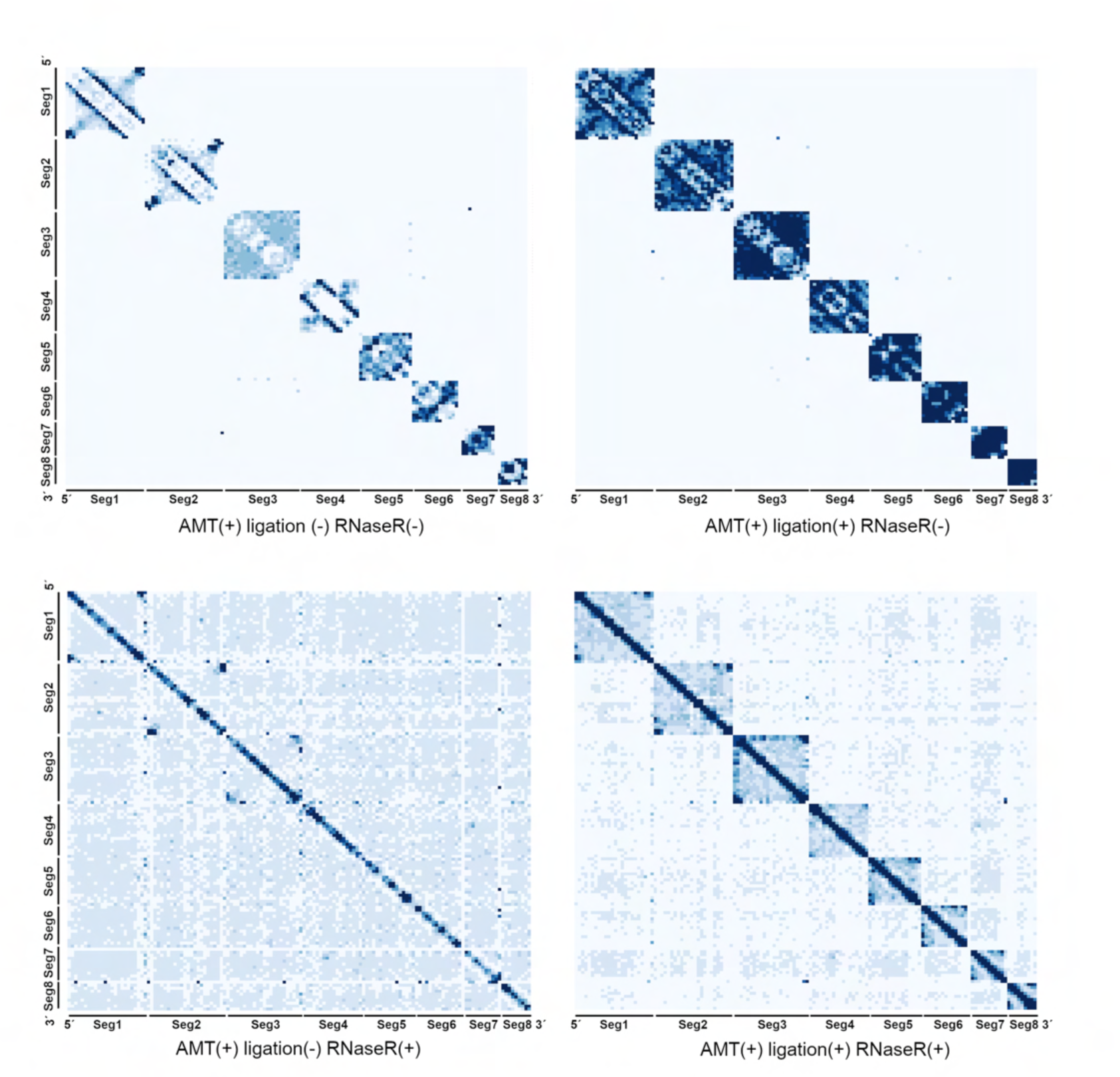
Control experiments of LIGR-seq. The normalized contact score matrix of control experiments of LIGR-seq was shown.

**Figure S10.**
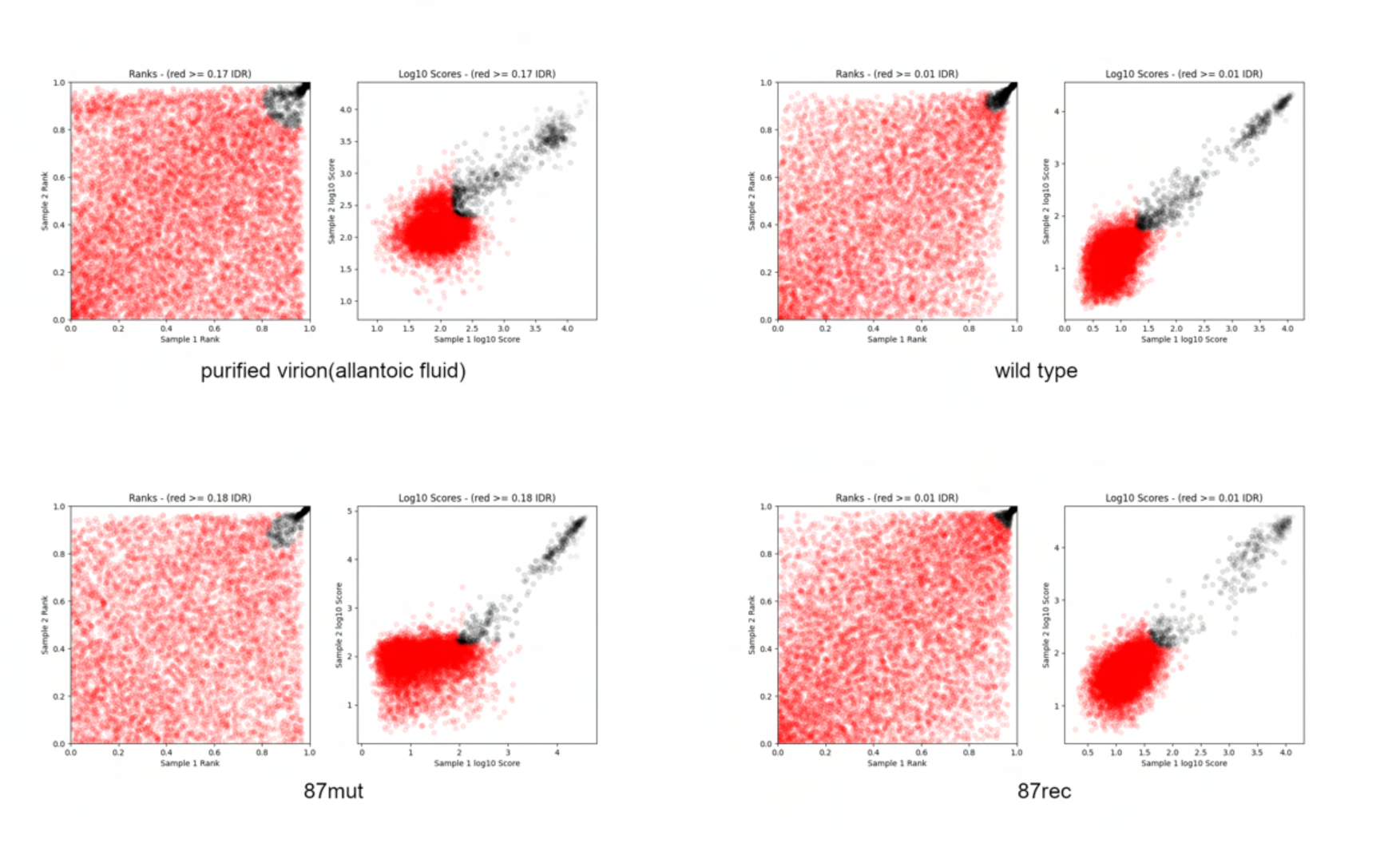
Reproducibility of LIGR-seq and identification of intersegment interactions with IDR scores from the top to the 100th. Duplicate LIGR-seq results from wild type (purified virion from allantoic fluid or cell culture supernatant), 87mut, and 87rec viruses were analyzed by IDR. Rank and log10 score of normalized contact score of each sample were plotted. The red circle in the purified virion from allantoic fluid, wild type, 87mut, 87rec virus panels indicates a region that IDR score is higher than threshold IDR score, and the black circle indicates a region that IDR score is lower than threshold IDR score. Threshold IDR score was determined by the IDR score of the 100th intersegment interaction and those of purified virion from allantoic fluid, wild type, 87mut, and 87rec viruses was 0.17, 0.01, 0.18, and 0.01, respectively. Red and black circles contain both intrasegment and intersegment interaction regions.

**Figure S11.**
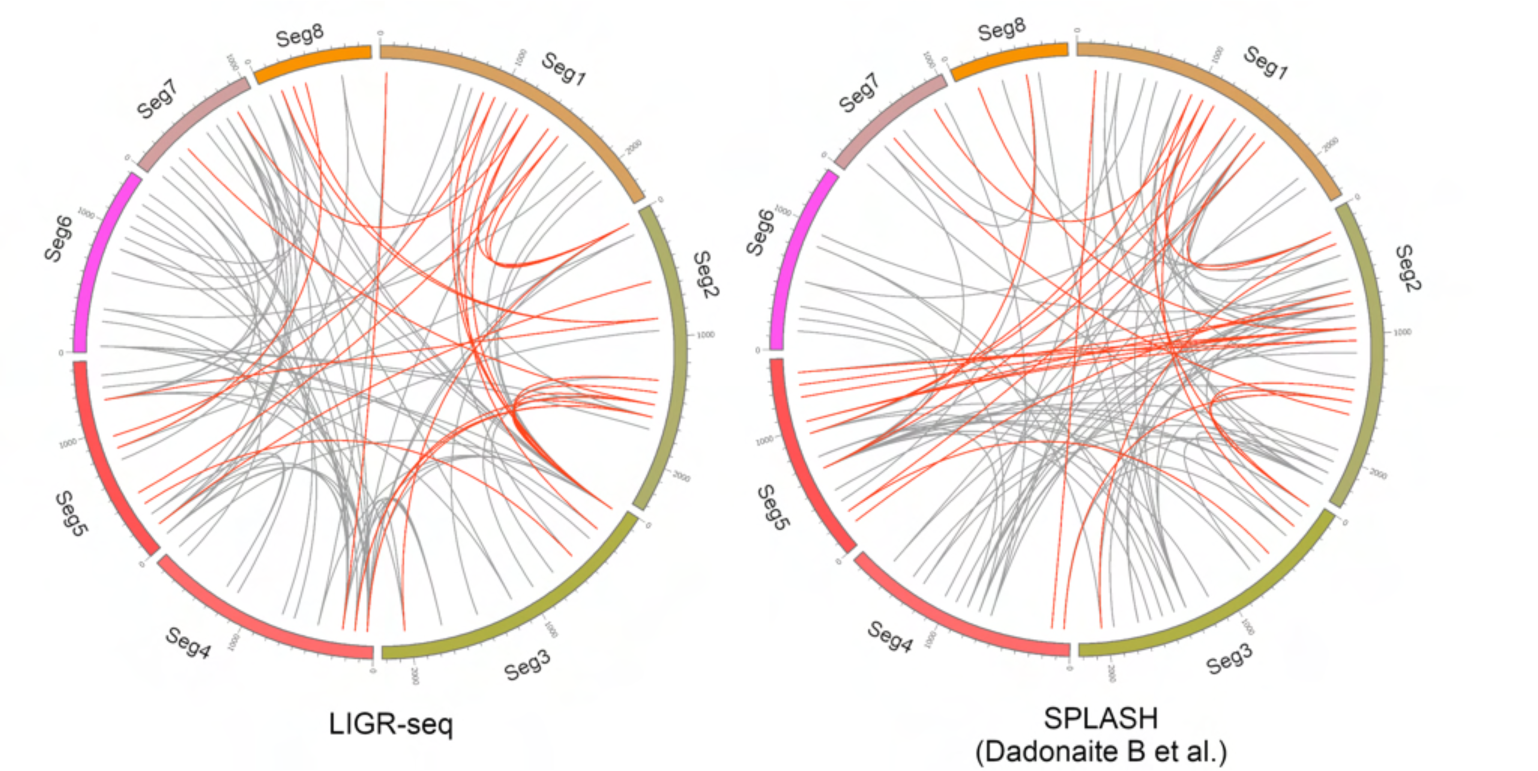
Comparison of the intersegment interactions identified by LIGR-seq with those by SPLASH. The intersegment interaction map of 100 interactions from LIGR-seq (left panel) and SPLASH (Dadonaite et al., 2019) (right panel). Intersegment interactions identified in both LIGR-seq and SPLASH within 200 nt were indicated by red lines.

**Figure S12.**
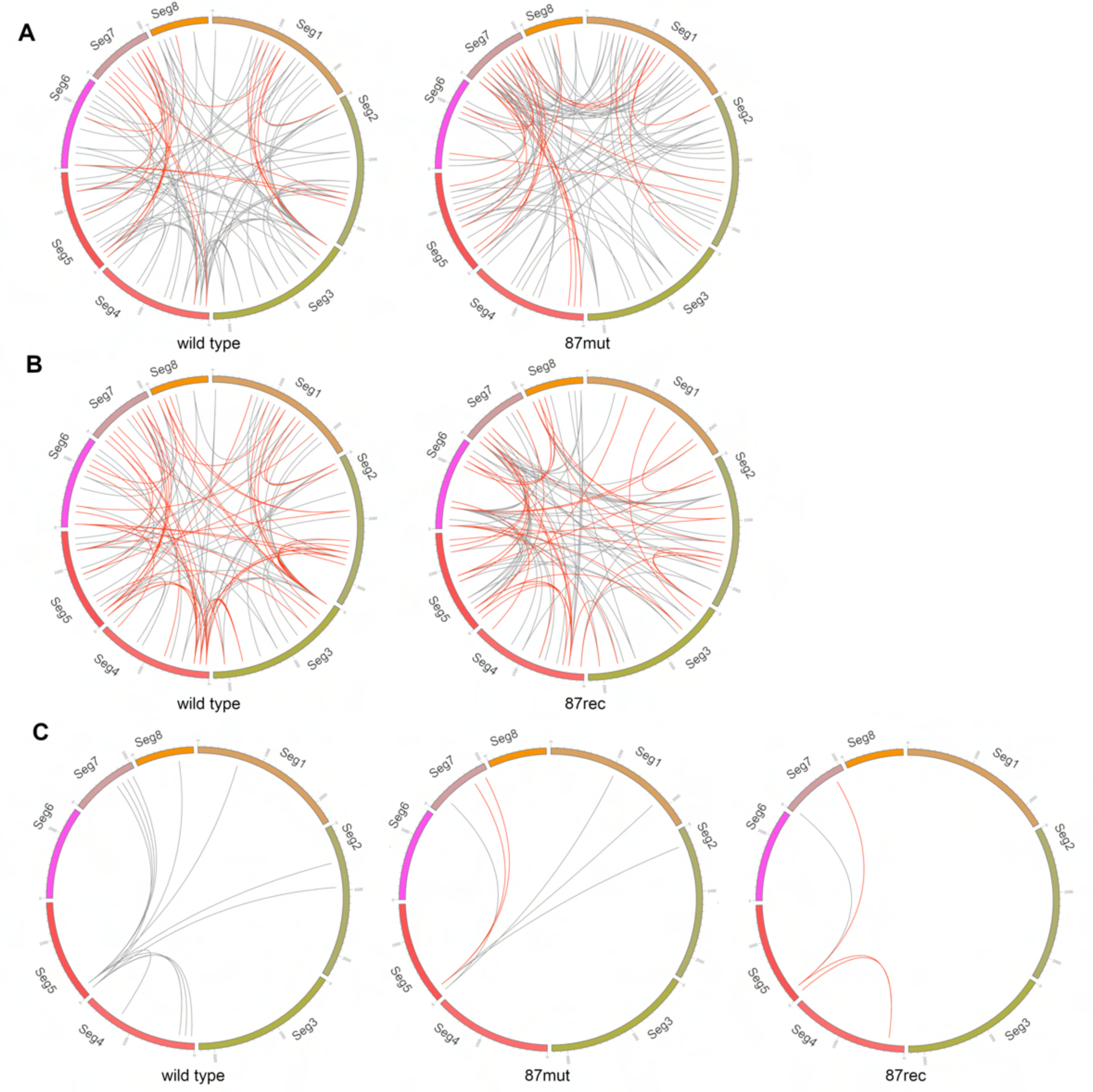
Comparison of the intersegment interactions among wild type, 87mut, and 87rec viruses. (A and B) Intersegment interactions between the wild type and 87mut (A) and wild type and 87rec viruses (B). Intersegment interaction maps of the wild type and 87mut virus (A) or the wild type and 87rec virus (B) were constructed from LIGR-seq. Intersegment interactions identified in both the wild type and the 87mut virus or the 87rec virus within a limit of 200 nt have been indicated by red lines. (C) The intersegment interactions between segment 5 and other segments in wild type, 87mut, and 87rec viruses. Intersegment interactions identified in both wild type virus and 87mut or 87rec viruses within 200 nt were indicated by red lines. Arrow indicates nucleotide positions 87-130 of segment 5 that form a pseudoknot structure.

**Table S1.**
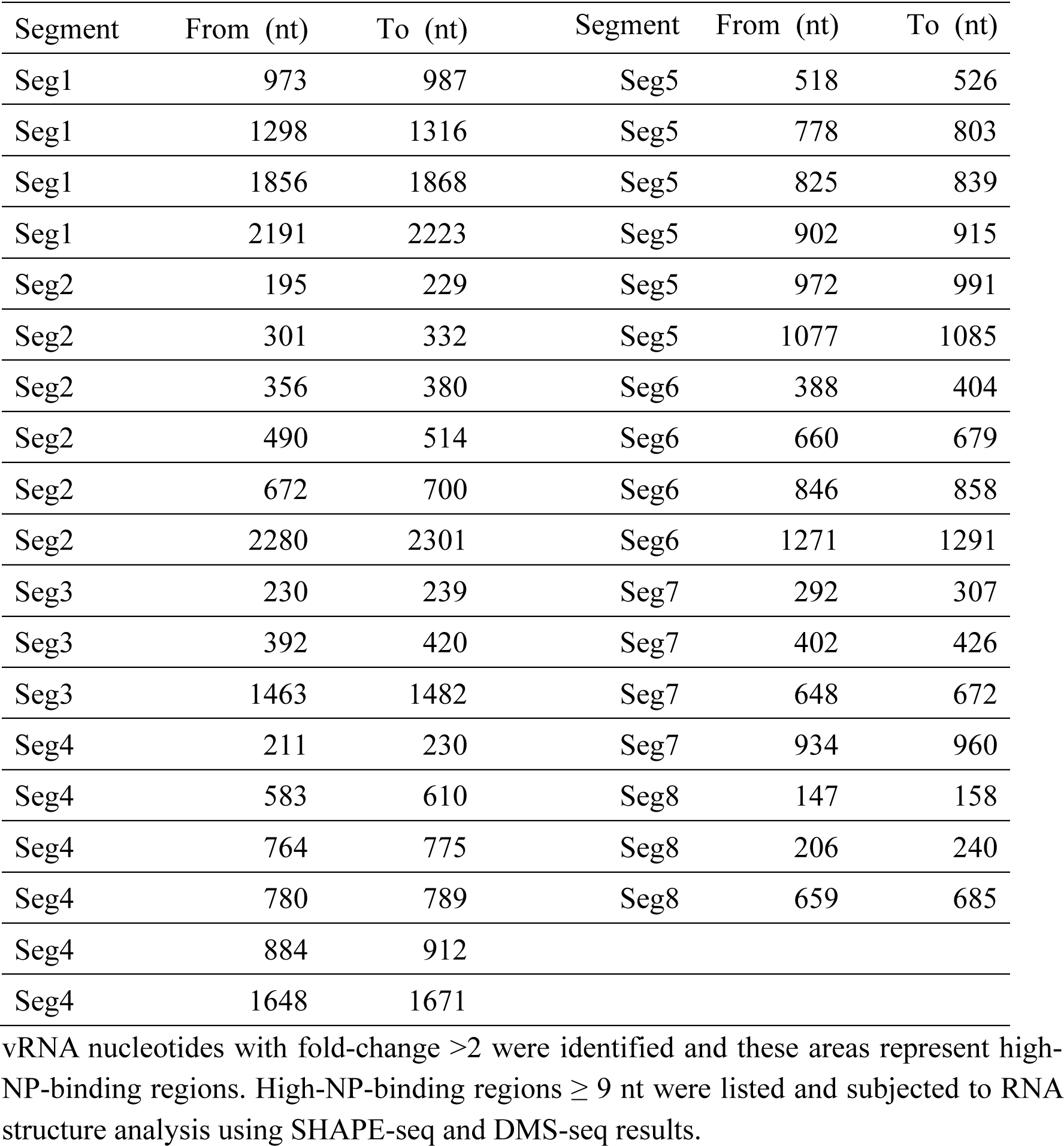
High-NP-binding regions from PAR-CLIP data sets

**Table S2.**
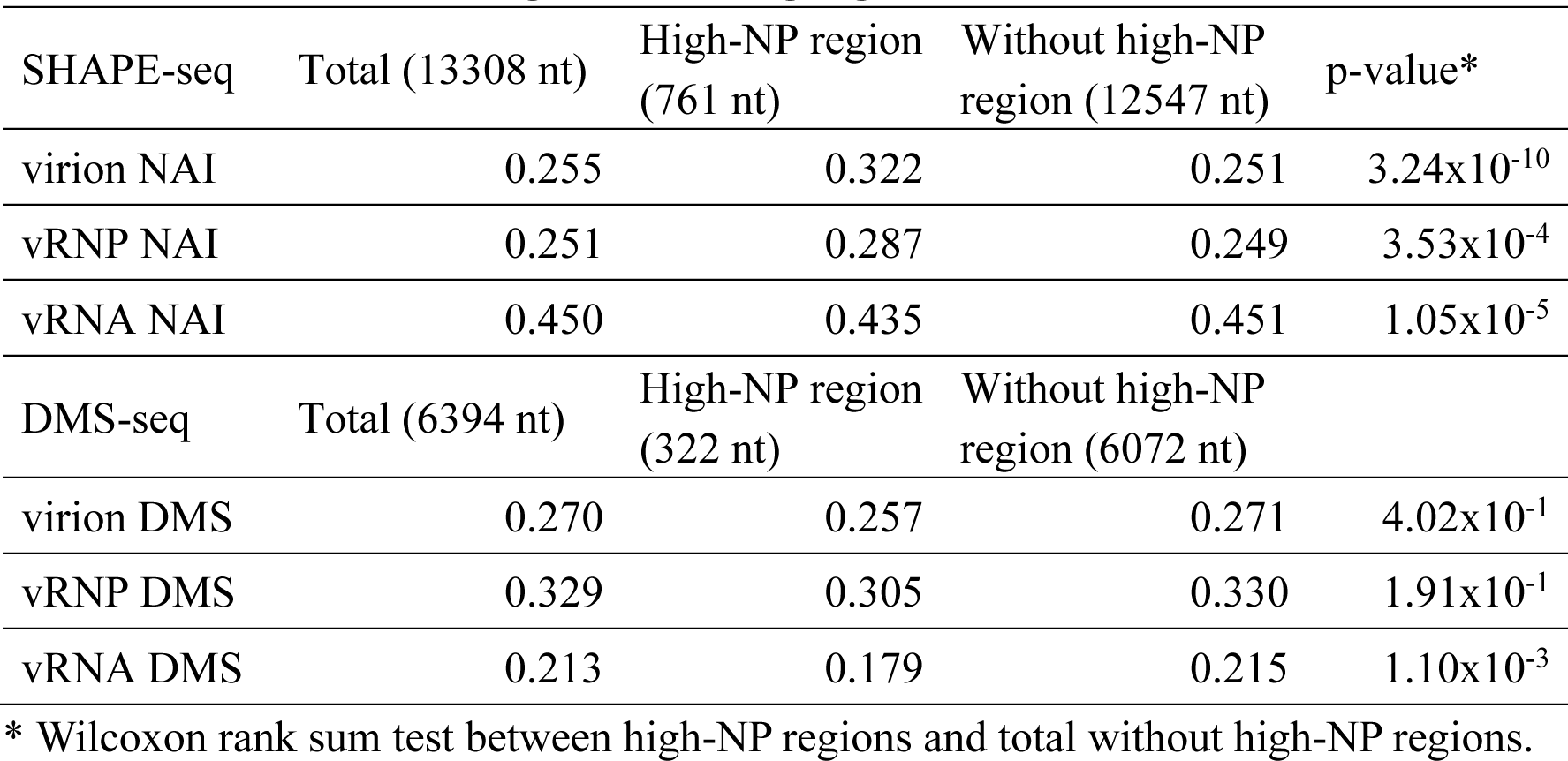
Probabilities of high-NP-binding regions

| SHAPE-seq | Total (13308 nt) | High-NP region (761 nt) | Without high-NP region (12547 nt) | p-value* |
| --- | --- | --- | --- | --- |
| virion NAI | 0.255 | 0.322 | 0.251 | $3.24 \times 10^{-10}$ |
| vRNP NAI | 0.251 | 0.287 | 0.249 | $3.53 \times 10^{-4}$ |
| vRNA NAI | 0.450 | 0.435 | 0.451 | $1.05 \times 10^{-5}$ |
| DMS-seq | Total (6394 nt) | High-NP region (322 nt) | Without high-NP region (6072 nt) |  |
| virion DMS | 0.270 | 0.257 | 0.271 | $4.02 \times 10^{-1}$ |
| vRNP DMS | 0.329 | 0.305 | 0.330 | $1.91 \times 10^{-1}$ |
| vRNA DMS | 0.213 | 0.179 | 0.215 | $1.10 \times 10^{-3}$ |

\* Wilcoxon rank sum test between high-NP regions and total without high-NP regions.

**Table S3.**
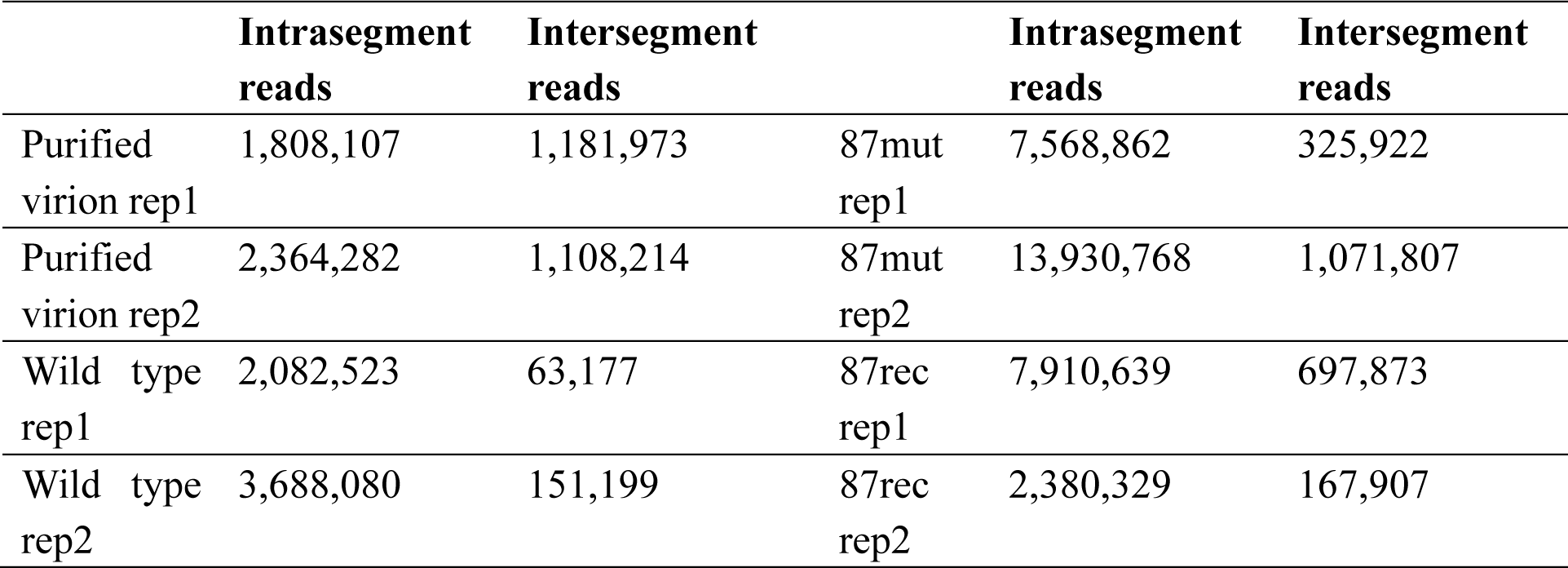
Intrasegment and intersegment mapped reads in LIGR-seq experiments.

## Notes

### Competing Interest Statement

The authors have declared no competing interest.

