## Supplementary material for "Comprehensive in virio structure probing analysis of the influenza A virus identifies a functional RNA structure involved in replication and segment interactions": Table S1

**Table S1. High-NP-binding regions from PAR-CLIP data sets**

| Segment | From (nt) | To (nt) |  | Segment | From (nt) | To (nt) |
| --- | --- | --- | --- | --- | --- | --- |
| Seg1 | 973 | 987 |  | Seg5 | 518 | 526 |
| Seg1 | 1298 | 1316 |  | Seg5 | 778 | 803 |
| Seg1 | 1856 | 1868 |  | Seg5 | 825 | 839 |
| Seg1 | 2191 | 2223 |  | Seg5 | 902 | 915 |
| Seg2 | 195 | 229 |  | Seg5 | 972 | 991 |
| Seg2 | 301 | 332 |  | Seg5 | 1077 | 1085 |
| Seg2 | 356 | 380 |  | Seg6 | 388 | 404 |
| Seg2 | 490 | 514 |  | Seg6 | 660 | 679 |
| Seg2 | 672 | 700 |  | Seg6 | 846 | 858 |
| Seg2 | 2280 | 2301 |  | Seg6 | 1271 | 1291 |
| Seg3 | 230 | 239 |  | Seg7 | 292 | 307 |
| Seg3 | 392 | 420 |  | Seg7 | 402 | 426 |
| Seg3 | 1463 | 1482 |  | Seg7 | 648 | 672 |
| Seg4 | 211 | 230 |  | Seg7 | 934 | 960 |
| Seg4 | 583 | 610 |  | Seg8 | 147 | 158 |
| Seg4 | 764 | 775 |  | Seg8 | 206 | 240 |
| Seg4 | 780 | 789 |  | Seg8 | 659 | 685 |
| Seg4 | 884 | 912 |  |  |  |  |
| Seg4 | 1648 | 1671 |  |  |  |  |

vRNA nucleotides with fold-change >2 were identified and these areas represent high-NP-binding regions. High-NP-binding regions ≥ 9 nt were listed and subjected to RNA structure analysis using SHAPE-seq and DMS-seq results.
