## Supplementary material for "Comprehensive in virio structure probing analysis of the influenza A virus identifies a functional RNA structure involved in replication and segment interactions": Table S2

**Table S2. Probabilities of high-NP-binding regions**

| SHAPE-seq | Total (13308 nt) | High-NP region (761 nt) | Without high-NP region (12547 nt) | p-value* |
| --- | --- | --- | --- | --- |
| virion NAI | 0.255 | 0.322 | 0.251 | 3.24x10^-10^ |
| vRNP NAI | 0.251 | 0.287 | 0.249 | 3.53x10^-4^ |
| vRNA NAI | 0.450 | 0.435 | 0.451 | 1.05x10^-5^ |
| DMS-seq | Total (6394 nt) | High-NP region (322 nt) | Without high-NP region (6072 nt) |  |
| virion DMS | 0.270 | 0.257 | 0.271 | 4.02x10^-1^ |
| vRNP DMS | 0.329 | 0.305 | 0.330 | 1.91x10^-1^ |
| vRNA DMS | 0.213 | 0.179 | 0.215 | 1.10x10^-3^ |

* Wilcoxon rank sum test between high-NP regions and total without high-NP regions.
