## Supplementary material for "Comprehensive in virio structure probing analysis of the influenza A virus identifies a functional RNA structure involved in replication and segment interactions": Table S3

**Table S3. Intrasegment and intersegment mapped reads in LIGR-seq experiments.**

|  | **Intrasegment reads** | **Intersegment reads** |  |  | **Intrasegment reads** | **Intersegment reads** |
| --- | --- | --- | --- | --- | --- | --- |
| Purified virion rep1 | 1,808,107 | 1,181,973 |  | 87mut rep1 | 7,568,862 | 325,922 |
| Purified virion rep2 | 2,364,282 | 1,108,214 |  | 87mut rep2 | 13,930,768 | 1,071,807 |
| Wild type rep1 | 2,082,523 | 63,177 |  | 87rec rep1 | 7,910,639 | 697,873 |
| Wild type rep2 | 3,688,080 | 151,199 |  | 87rec rep2 | 2,380,329 | 167,907 |
