## Supplementary material for "Comprehensive in virio structure probing analysis of the influenza A virus identifies a functional RNA structure involved in replication and segment interactions": Table S4

**Table S4. Primers used in this study**

| Recombinant virus preparation | |
| --- | --- |
| Name | **Sequence** |
| PR8_Seg5_87_130mut_rev | GCAGCGAGCCCGATTGTGCCTTCC |
| PR8_Seg5_87_130mut_for | CTTTTCGTCCGAGAGTTCAAAGAC |
| PR8_Seg5_87_130rec_rev | GCAGCAAGTCCAATTGTGCCTTCC |
| RT-qPCR | |
| Name | **Sequence** |
| Uni12 | AGCRAAAGCAGG |
| Seg1_qPCR_for | AGCAAGCCGTGGATATTTGC |
| Seg1_qPCR_rev | TGAAGATTGCCCGTAAGCAC |
| Seg2_qPCR_for | TGTTCAGCAAACACGAGTGG |
| Seg2_qPCR_rev | TGTGTTGGCCAATGCTGTTG |
| Seg3_qPCR_for | TTGAGGTCGCTTGCAAGTTG |
| Seg3_qPCR_rev | TGTTCAATTGGAGCCGCATC |
| Seg4_qPCR_for | GGCATCATCACCTCAAACGCGTC |
| Seg4_qPCR_rev | ATCCTCAATTTGGTACTCCTGACA |
| Seg5_qPCR_for | TAATGGTGACGATGCAACGG |
| Seg5_qPCR_rev | ATTCCTGTGCGAACAAGAGC |
| Seg6_qPCR_for | ACAATTGGCACGGTTCGAAC |
| Seg6_qPCR_rev | AGCTGCCTGTTCCATCTTTG |
| Seg7_qPCR_for | AGAGGGAGATAACATTCCATGGGGC |
| Seg7_qPCR_rev | TGTTCACAGGTTGCGCATACCAGGC |
| Seg8_qPCR_2_for | TCCAGCCGGTCAAAAATCAC |
| Seg8_qPCR_2_rev | TGAGGAAATGTCAAGGGACTGG |
| DMS-seq and SHAPE-seq | |
| Name | **Sequence** |
| Random_hexamer_adaptor | CAGACGTGTGCTCTTCCGATCTNNNNNN |
| ssDNA_linker | /5Phos/NNNAGATCGGAAGAGCGTCGTGTAG/3SpC3/ |
| illumina_TurSeq_forward | AATGATACGGCGACCACCGAGATCTACACTCTTTCCCTACACGACGCTCTTCCGATCT |
| illumina_barcode1_PCR | CAAGCAGAAGACGGCATACGAGATCGTGATGTGACTGGAGTTCAGACGTGTGCTCTTCCGATCT |
| illumina_barcode2_PCR | CAAGCAGAAGACGGCATACGAGATACATCGGTGACTGGAGTTCAGACGTGTGCTCTTCCGATCT |
| illumina_barcode3_PCR | CAAGCAGAAGACGGCATACGAGATGCCTAAGTGACTGGAGTTCAGACGTGTGCTCTTCCGATCT |
| illumina_barcode4_PCR | CAAGCAGAAGACGGCATACGAGATTGGTCAGTGACTGGAGTTCAGACGTGTGCTCTTCCGATCT |
| illumina_barcode5_PCR | CAAGCAGAAGACGGCATACGAGATCACTGTGTGACTGGAGTTCAGACGTGTGCTCTTCCGATCT |
| illumina_barcode6_PCR | CAAGCAGAAGACGGCATACGAGATATTGGCGTGACTGGAGTTCAGACGTGTGCTCTTCCGATCT |
| illumina_barcode7_PCR | CAAGCAGAAGACGGCATACGAGATGATCTGGTGACTGGAGTTCAGACGTGTGCTCTTCCGATCT |
| illumina_barcode8_PCR | CAAGCAGAAGACGGCATACGAGATTCAAGTGTGACTGGAGTTCAGACGTGTGCTCTTCCGATCT |
| illumina_barcode9_PCR | CAAGCAGAAGACGGCATACGAGATCTGATCGTGACTGGAGTTCAGACGTGTGCTCTTCCGATCT |
| illumina_barcode10_PCR | CAAGCAGAAGACGGCATACGAGATAAGCTAGTGACTGGAGTTCAGACGTGTGCTCTTCCGATCT |
| illumina_barcode11_PCR | CAAGCAGAAGACGGCATACGAGATGTAGCCGTGACTGGAGTTCAGACGTGTGCTCTTCCGATCT |
| illumina_barcode12_PCR | CAAGCAGAAGACGGCATACGAGATTACAAGGTGACTGGAGTTCAGACGTGTGCTCTTCCGATCT |
| RNA pull-down | |
| Name | **Sequence** |
| Biotin_Linker1 for C probe | GTCAGTCAGTCAGTCAGTCA |
| Linker2 | GGATCCAATTGGCCAACTCG |
| S1_Linker1_28_for | GTCAGTCAGTCAGTCAGTCAATGGAAAGAATAAAAGAACT |
| S1_Linker1_133_for | GTCAGTCAGTCAGTCAGTCATCAGGAAGACAGGAGAAGAA |
| S1_Linker1_238_for | GTCAGTCAGTCAGTCAGTCAAATGAGCAAGGACAAACTTT |
| S1_Linker1_343_for | GTCAGTCAGTCAGTCAGTCAACAAATACAGTTCATTATCC |
| S1_Linker1_448_for | GTCAGTCAGTCAGTCAGTCAATACGTCGGAGAGTTGACAT |
| S1_Linker1_553_for | GTCAGTCAGTCAGTCAGTCAATACTAACATCGGAATCGCA |
| S1_Linker1_658_for | GTCAGTCAGTCAGTCAGTCACTGGTCCGCAAAACGAGATT |
| S1_Linker1_763_for | GTCAGTCAGTCAGTCAGTCACCAGGAGGGGAAGTGAGGAA |
| S1_Linker1_868_for | GTCAGTCAGTCAGTCAGTCACTGGAGATGTGCCACAGCAC |
| S1_Linker1_973_for | GTCAGTCAGTCAGTCAGTCAGGACTGAGAATTAGCTCATC |
| S1_Linker1_1078_for | GTCAGTCAGTCAGTCAGTCAACATTGAAGATAGGAGTGCA |
| S1_Linker1_1183_for | GTCAGTCAGTCAGTCAGTCAGTGAGTGGGAGAGACGAACA |
| S1_Linker1_1288_for | GTCAGTCAGTCAGTCAGTCAGTCAATAGGGCGAATCAGCG |
| S1_Linker1_1393_for | GTCAGTCAGTCAGTCAGTCAAATGTGATGGGAATGATTGG |
| S1_Linker1_1498_for | GTCAGTCAGTCAGTCAGTCAACGGAGAGGGTAGTGGTGAG |
| S1_Linker1_1603_for | GTCAGTCAGTCAGTCAGTCAAAACTGACAATAACTTACTC |
| S1_Linker1_1708_for | GTCAGTCAGTCAGTCAGTCAAAAATTCAGTGGTCCCAGAA |
| S1_Linker1_1813_for | GTCAGTCAGTCAGTCAGTCAGTAAGAACTCTGTTCCAACA |
| S1_Linker1_1918_for | GTCAGTCAGTCAGTCAGTCAATGCAGTTCTCCTCATTTAC |
| S1_Linker1_2023_for | GTCAGTCAGTCAGTCAGTCAACAGTTCTCGGAAAGGATGC |
| S1_Linker1_2128_for | GTCAGTCAGTCAGTCAGTCAGACAAGAGATATGGGCCAGC |
| S1_Linker2_147_rev | GGATCCAATTGGCCAACTCGCTCCTGTCTTCCTGATGTGT |
| S1_Linker2_252_rev | GGATCCAATTGGCCAACTCGTTGTCCTTGCTCATTTCTCT |
| S1_Linker2_357_rev | GGATCCAATTGGCCAACTCGATGAACTGTATTTGTTATTG |
| S1_Linker2_462_rev | GGATCCAATTGGCCAACTCGAACTCTCCGACGTATTTTGA |
| S1_Linker2_567_rev | GGATCCAATTGGCCAACTCGTTCCGATGTTAGTATCCTGG |
| S1_Linker2_672_rev | GGATCCAATTGGCCAACTCGCGTTTTGCGGACCAGTTCTC |
| S1_Linker2_777_rev | GGATCCAATTGGCCAACTCGCACTTCCCCTCCTGGAGTAT |
| S1_Linker2_882_rev | GGATCCAATTGGCCAACTCGGTGGCACATCTCCAGTAAAG |
| S1_Linker2_987_rev | GGATCCAATTGGCCAACTCGGCTAATTCTCAGTCCCATTG |
| S1_Linker2_1092_rev | GGATCCAATTGGCCAACTCGTCCTATCTTCAATGTTTGAA |
| S1_Linker2_1197_rev | GGATCCAATTGGCCAACTCGGTCTCTCCCACTCACTATCA |
| S1_Linker2_1302_rev | GGATCCAATTGGCCAACTCGATTCGCCCTATTGACGAAAT |
| S1_Linker2_1407_rev | GGATCCAATTGGCCAACTCGCATTCCCATCACATTGTCGA |
| S1_Linker2_1512_rev | GGATCCAATTGGCCAACTCGCACTACCCTCTCCGTGCTGG |
| S1_Linker2_1617_rev | GGATCCAATTGGCCAACTCGAGTTATTGTCAGTTTCTCTG |
| S1_Linker2_1722_rev | GGATCCAATTGGCCAACTCGGGACCACTGAATTTTAACAG |
| S1_Linker2_1827_rev | GGATCCAATTGGCCAACTCGGAACAGAGTTCTTACAAACC |
| S1_Linker2_1932_rev | GGATCCAATTGGCCAACTCGTGAGGAGAACTGCATTCTAC |
| S1_Linker2_2037_rev | GGATCCAATTGGCCAACTCGCTTTCCGAGAACTGTGAGTC |
| S1_Linker2_2142_rev | GGATCCAATTGGCCAACTCGCCCATATCTCTTGTCTTCTT |
| S1_Linker2_2247_rev | GGATCCAATTGGCCAACTCGGTCCCGTTTCCGTTTCATTA |
| S2_Linker1_25_for | GTCAGTCAGTCAGTCAGTCAATGGATGTCAATCCGACCTT |
| S2_Linker1_130_for | GTCAGTCAGTCAGTCAGTCAACAGGATACACCATGGATAC |
| S2_Linker1_235_for | GTCAGTCAGTCAGTCAGTCAGGGCCACTGCCAGAAGACAA |
| S2_Linker1_340_for | GTCAGTCAGTCAGTCAGTCATCGTGTATTGAAACGATGGA |
| S2_Linker1_445_for | GTCAGTCAGTCAGTCAGTCAACAGCATTGGCCAACACAAT |
| S2_Linker1_550_for | GTCAGTCAGTCAGTCAGTCAAAAGAAGAAATGGGGATCAC |
| S2_Linker1_655_for | GTCAGTCAGTCAGTCAGTCAAGATTGAACAAAAGGAGTTA |
| S2_Linker1_760_for | GTCAGTCAGTCAGTCAGTCAATGCAAATAAGGGGGTTTGT |
| S2_Linker1_865_for | GTCAGTCAGTCAGTCAGTCAAAGTTGGCAAATGTTGTAAG |
| S2_Linker1_970_for | GTCAGTCAGTCAGTCAGTCACGGATGTTTTTGGCCATGAT |
| S2_Linker1_1075_for | GTCAGTCAGTCAGTCAGTCACTGGGAAAAGGGTATATGTT |
| S2_Linker1_1180_for | GTCAGTCAGTCAGTCAGTCAAGAAAGAAGATTGAAAAAAT |
| S2_Linker1_1285_for | GTCAGTCAGTCAGTCAGTCAGTCTCCATCCTGAATCTTGG |
| S2_Linker1_1390_for | GTCAGTCAGTCAGTCAGTCACATGAAGGGATTCAAGCCGG |
| S2_Linker1_1495_for | GTCAGTCAGTCAGTCAGTCAGAATTCACAAGTTTTTTCTA |
| S2_Linker1_1600_for | GTCAGTCAGTCAGTCAGTCAGGAGTTACTGTCATCAAAAA |
| S2_Linker1_1705_for | GTCAGTCAGTCAGTCAGTCATGCCATAGAGGTGACACACA |
| S2_Linker1_1810_for | GTCAGTCAGTCAGTCAGTCACCAAATTTATACAACATTAG |
| S2_Linker1_1915_for | GTCAGTCAGTCAGTCAGTCATTTGTCAGCCATAAAGAAAT |
| S2_Linker1_2020_for | GTCAGTCAGTCAGTCAGTCATGGATCCCCAAAAGAAATCG |
| S2_Linker1_2125_for | GTCAGTCAGTCAGTCAGTCACCCAGCAGTTCATACAGAAG |
| S2_Linker2_144_rev | GGATCCAATTGGCCAACTCGCATGGTGTATCCTGTTCCTG |
| S2_Linker2_249_rev | GGATCCAATTGGCCAACTCGTTCTGGCAGTGGCCCATCAA |
| S2_Linker2_354_rev | GGATCCAATTGGCCAACTCGCGTTTCAATACACGAGTTTT |
| S2_Linker2_459_rev | GGATCCAATTGGCCAACTCGGTTGGCCAATGCTGTTGCAG |
| S2_Linker2_564_rev | GGATCCAATTGGCCAACTCGCCCCATTTCTTCTTTGTTCA |
| S2_Linker2_669_rev | GGATCCAATTGGCCAACTCGCCTTTTGTTCAATCTCTGCT |
| S2_Linker2_774_rev | GGATCCAATTGGCCAACTCGCCCCCTTATTTGCATCCCTG |
| S2_Linker2_879_rev | GGATCCAATTGGCCAACTCGAACATTTGCCAACTTTGCTT |
| S2_Linker2_984_rev | GGATCCAATTGGCCAACTCGGGCCAAAAACATCCGAGGAT |
| S2_Linker2_1089_rev | GGATCCAATTGGCCAACTCGATACCCTTTTCCCAGTCTCG |
| S2_Linker2_1194_rev | GGATCCAATTGGCCAACTCGTTCAATCTTCTTTCTTGTTG |
| S2_Linker2_1299_rev | GGATCCAATTGGCCAACTCGATTCAGGATGGAGACGCCTA |
| S2_Linker2_1404_rev | GGATCCAATTGGCCAACTCGTTGAATCCCTTCATGATTGG |
| S2_Linker2_1509_rev | GGATCCAATTGGCCAACTCGAAAACTTGTGAATTCAAATG |
| S2_Linker2_1614_rev | GGATCCAATTGGCCAACTCGGATGACAGTAACTCCAATAC |
| S2_Linker2_1719_rev | GGATCCAATTGGCCAACTCGGTCACCTCTATGGCATCGGT |
| S2_Linker2_1824_rev | GGATCCAATTGGCCAACTCGGTTGTATAAATTTGGGCCTC |
| S2_Linker2_1929_rev | GGATCCAATTGGCCAACTCGTTTATGGCTGACAAATGGGT |
| S2_Linker2_2034_rev | GGATCCAATTGGCCAACTCGTCTTTTGGGGATCCAGGAGT |
| S2_Linker2_2139_rev | GGATCCAATTGGCCAACTCGGTATGAACTGCTGGGGAAGA |
| S2_Linker2_2244_rev | GGATCCAATTGGCCAACTCGGAACTCTTCTTTCTTTATCC |
| S1_Linker1_28_for | GTCAGTCAGTCAGTCAGTCAATGGAAAGAATAAAAGAACT |
| S1_Linker1_133_for | GTCAGTCAGTCAGTCAGTCATCAGGAAGACAGGAGAAGAA |
| S1_Linker1_238_for | GTCAGTCAGTCAGTCAGTCAAATGAGCAAGGACAAACTTT |
| S1_Linker1_343_for | GTCAGTCAGTCAGTCAGTCAACAAATACAGTTCATTATCC |
| S1_Linker1_448_for | GTCAGTCAGTCAGTCAGTCAATACGTCGGAGAGTTGACAT |
| S1_Linker1_553_for | GTCAGTCAGTCAGTCAGTCAATACTAACATCGGAATCGCA |
| S1_Linker1_658_for | GTCAGTCAGTCAGTCAGTCACTGGTCCGCAAAACGAGATT |
| S1_Linker1_763_for | GTCAGTCAGTCAGTCAGTCACCAGGAGGGGAAGTGAGGAA |
| S1_Linker1_868_for | GTCAGTCAGTCAGTCAGTCACTGGAGATGTGCCACAGCAC |
| S1_Linker1_973_for | GTCAGTCAGTCAGTCAGTCAGGACTGAGAATTAGCTCATC |
| S1_Linker1_1078_for | GTCAGTCAGTCAGTCAGTCAACATTGAAGATAGGAGTGCA |
| S1_Linker1_1183_for | GTCAGTCAGTCAGTCAGTCAGTGAGTGGGAGAGACGAACA |
| S1_Linker1_1288_for | GTCAGTCAGTCAGTCAGTCAGTCAATAGGGCGAATCAGCG |
| S1_Linker1_1393_for | GTCAGTCAGTCAGTCAGTCAAATGTGATGGGAATGATTGG |
| S1_Linker1_1498_for | GTCAGTCAGTCAGTCAGTCAACGGAGAGGGTAGTGGTGAG |
| S1_Linker1_1603_for | GTCAGTCAGTCAGTCAGTCAAAACTGACAATAACTTACTC |
| S1_Linker1_1708_for | GTCAGTCAGTCAGTCAGTCAAAAATTCAGTGGTCCCAGAA |
| S1_Linker1_1813_for | GTCAGTCAGTCAGTCAGTCAGTAAGAACTCTGTTCCAACA |
| S1_Linker1_1918_for | GTCAGTCAGTCAGTCAGTCAATGCAGTTCTCCTCATTTAC |
| S1_Linker1_2023_for | GTCAGTCAGTCAGTCAGTCAACAGTTCTCGGAAAGGATGC |
| S1_Linker1_2128_for | GTCAGTCAGTCAGTCAGTCAGACAAGAGATATGGGCCAGC |
| S1_Linker2_147_rev | GGATCCAATTGGCCAACTCGCTCCTGTCTTCCTGATGTGT |
| S1_Linker2_252_rev | GGATCCAATTGGCCAACTCGTTGTCCTTGCTCATTTCTCT |
| S1_Linker2_357_rev | GGATCCAATTGGCCAACTCGATGAACTGTATTTGTTATTG |
| S1_Linker2_462_rev | GGATCCAATTGGCCAACTCGAACTCTCCGACGTATTTTGA |
| S1_Linker2_567_rev | GGATCCAATTGGCCAACTCGTTCCGATGTTAGTATCCTGG |
| S1_Linker2_672_rev | GGATCCAATTGGCCAACTCGCGTTTTGCGGACCAGTTCTC |
| S1_Linker2_777_rev | GGATCCAATTGGCCAACTCGCACTTCCCCTCCTGGAGTAT |
| S1_Linker2_882_rev | GGATCCAATTGGCCAACTCGGTGGCACATCTCCAGTAAAG |
| S1_Linker2_987_rev | GGATCCAATTGGCCAACTCGGCTAATTCTCAGTCCCATTG |
| S1_Linker2_1092_rev | GGATCCAATTGGCCAACTCGTCCTATCTTCAATGTTTGAA |
| S1_Linker2_1197_rev | GGATCCAATTGGCCAACTCGGTCTCTCCCACTCACTATCA |
| S1_Linker2_1302_rev | GGATCCAATTGGCCAACTCGATTCGCCCTATTGACGAAAT |
| S1_Linker2_1407_rev | GGATCCAATTGGCCAACTCGCATTCCCATCACATTGTCGA |
| S1_Linker2_1512_rev | GGATCCAATTGGCCAACTCGCACTACCCTCTCCGTGCTGG |
| S1_Linker2_1617_rev | GGATCCAATTGGCCAACTCGAGTTATTGTCAGTTTCTCTG |
| S1_Linker2_1722_rev | GGATCCAATTGGCCAACTCGGGACCACTGAATTTTAACAG |
| S1_Linker2_1827_rev | GGATCCAATTGGCCAACTCGGAACAGAGTTCTTACAAACC |
| S1_Linker2_1932_rev | GGATCCAATTGGCCAACTCGTGAGGAGAACTGCATTCTAC |
| S1_Linker2_2037_rev | GGATCCAATTGGCCAACTCGCTTTCCGAGAACTGTGAGTC |
| S1_Linker2_2142_rev | GGATCCAATTGGCCAACTCGCCCATATCTCTTGTCTTCTT |
| S1_Linker2_2247_rev | GGATCCAATTGGCCAACTCGGTCCCGTTTCCGTTTCATTA |
| S2_Linker1_25_for | GTCAGTCAGTCAGTCAGTCAATGGATGTCAATCCGACCTT |
| S2_Linker1_130_for | GTCAGTCAGTCAGTCAGTCAACAGGATACACCATGGATAC |
| S2_Linker1_235_for | GTCAGTCAGTCAGTCAGTCAGGGCCACTGCCAGAAGACAA |
| S2_Linker1_340_for | GTCAGTCAGTCAGTCAGTCATCGTGTATTGAAACGATGGA |
| S2_Linker1_445_for | GTCAGTCAGTCAGTCAGTCAACAGCATTGGCCAACACAAT |
| S2_Linker1_550_for | GTCAGTCAGTCAGTCAGTCAAAAGAAGAAATGGGGATCAC |
| S2_Linker1_655_for | GTCAGTCAGTCAGTCAGTCAAGATTGAACAAAAGGAGTTA |
| S2_Linker1_760_for | GTCAGTCAGTCAGTCAGTCAATGCAAATAAGGGGGTTTGT |
| S2_Linker1_865_for | GTCAGTCAGTCAGTCAGTCAAAGTTGGCAAATGTTGTAAG |
| S2_Linker1_970_for | GTCAGTCAGTCAGTCAGTCACGGATGTTTTTGGCCATGAT |
| S2_Linker1_1075_for | GTCAGTCAGTCAGTCAGTCACTGGGAAAAGGGTATATGTT |
| S2_Linker1_1180_for | GTCAGTCAGTCAGTCAGTCAAGAAAGAAGATTGAAAAAAT |
| S2_Linker1_1285_for | GTCAGTCAGTCAGTCAGTCAGTCTCCATCCTGAATCTTGG |
| S2_Linker1_1390_for | GTCAGTCAGTCAGTCAGTCACATGAAGGGATTCAAGCCGG |
| S2_Linker1_1495_for | GTCAGTCAGTCAGTCAGTCAGAATTCACAAGTTTTTTCTA |
| S2_Linker1_1600_for | GTCAGTCAGTCAGTCAGTCAGGAGTTACTGTCATCAAAAA |
| S2_Linker1_1705_for | GTCAGTCAGTCAGTCAGTCATGCCATAGAGGTGACACACA |
| S2_Linker1_1810_for | GTCAGTCAGTCAGTCAGTCACCAAATTTATACAACATTAG |
| S2_Linker1_1915_for | GTCAGTCAGTCAGTCAGTCATTTGTCAGCCATAAAGAAAT |
| S2_Linker1_2020_for | GTCAGTCAGTCAGTCAGTCATGGATCCCCAAAAGAAATCG |
| S2_Linker1_2125_for | GTCAGTCAGTCAGTCAGTCACCCAGCAGTTCATACAGAAG |
| S2_Linker2_144_rev | GGATCCAATTGGCCAACTCGCATGGTGTATCCTGTTCCTG |
| S2_Linker2_249_rev | GGATCCAATTGGCCAACTCGTTCTGGCAGTGGCCCATCAA |
| S2_Linker2_354_rev | GGATCCAATTGGCCAACTCGCGTTTCAATACACGAGTTTT |
| S2_Linker2_459_rev | GGATCCAATTGGCCAACTCGGTTGGCCAATGCTGTTGCAG |
| S2_Linker2_564_rev | GGATCCAATTGGCCAACTCGCCCCATTTCTTCTTTGTTCA |
| S2_Linker2_669_rev | GGATCCAATTGGCCAACTCGCCTTTTGTTCAATCTCTGCT |
| S2_Linker2_774_rev | GGATCCAATTGGCCAACTCGCCCCCTTATTTGCATCCCTG |
| S2_Linker2_879_rev | GGATCCAATTGGCCAACTCGAACATTTGCCAACTTTGCTT |
| S2_Linker2_984_rev | GGATCCAATTGGCCAACTCGGGCCAAAAACATCCGAGGAT |
| S2_Linker2_1089_rev | GGATCCAATTGGCCAACTCGATACCCTTTTCCCAGTCTCG |
| S2_Linker2_1194_rev | GGATCCAATTGGCCAACTCGTTCAATCTTCTTTCTTGTTG |
| S2_Linker2_1299_rev | GGATCCAATTGGCCAACTCGATTCAGGATGGAGACGCCTA |
| S2_Linker2_1404_rev | GGATCCAATTGGCCAACTCGTTGAATCCCTTCATGATTGG |
| S2_Linker2_1509_rev | GGATCCAATTGGCCAACTCGAAAACTTGTGAATTCAAATG |
| S2_Linker2_1614_rev | GGATCCAATTGGCCAACTCGGATGACAGTAACTCCAATAC |
| S2_Linker2_1719_rev | GGATCCAATTGGCCAACTCGGTCACCTCTATGGCATCGGT |
| S2_Linker2_1824_rev | GGATCCAATTGGCCAACTCGGTTGTATAAATTTGGGCCTC |
| S2_Linker2_1929_rev | GGATCCAATTGGCCAACTCGTTTATGGCTGACAAATGGGT |
| S2_Linker2_2034_rev | GGATCCAATTGGCCAACTCGTCTTTTGGGGATCCAGGAGT |
| S2_Linker2_2139_rev | GGATCCAATTGGCCAACTCGGTATGAACTGCTGGGGAAGA |
| S2_Linker2_2244_rev | GGATCCAATTGGCCAACTCGGAACTCTTCTTTCTTTATCC |
| S3_Linker1_25_for | GTCAGTCAGTCAGTCAGTCAATGGAAGATTTTGTGCGACA |
| S3_Linker1_130_for | GTCAGTCAGTCAGTCAGTCAGCAGCAATATGCACTCACTT |
| S3_Linker1_235_for | GTCAGTCAGTCAGTCAGTCACTTTTGAAGCACAGATTTGA |
| S3_Linker1_340_for | GTCAGTCAGTCAGTCAGTCACTACCAGATTTGTATGATTA |
| S3_Linker1_445_for | GTCAGTCAGTCAGTCAGTCAGAGGAAACACACATCCACAT |
| S3_Linker1_550_for | GTCAGTCAGTCAGTCAGTCATTCACTATAAGACAAGAAAT |
| S3_Linker1_655_for | GTCAGTCAGTCAGTCAGTCAATGCGCAAGCTTGCCGACCA |
| S3_Linker1_760_for | GTCAGTCAGTCAGTCAGTCACTGTCTCAAATGTCCAAAGA |
| S3_Linker1_865_for | GTCAGTCAGTCAGTCAGTCAAAATTCCTGCTGATGGATGC |
| S3_Linker1_970_for | GTCAGTCAGTCAGTCAGTCAGGATGGAAGGAACCCAATGT |
| S3_Linker1_1075_for | GTCAGTCAGTCAGTCAGTCAGAGGAGAAAATTCCAAAGAC |
| S3_Linker1_1180_for | GTCAGTCAGTCAGTCAGTCAGATGTAGGTGATTTGAAGCA |
| S3_Linker1_1285_for | GTCAGTCAGTCAGTCAGTCAAGCTGGATAGAGCTCGATGA |
| S3_Linker1_1390_for | GTCAGTCAGTCAGTCAGTCAACAGAATACATAATGAAGGG |
| S3_Linker1_1495_for | GTCAGTCAGTCAGTCAGTCAACTAAGGAGGGAAGGCGAAA |
| S3_Linker1_1600_for | GTCAGTCAGTCAGTCAGTCATCTCTCACTGACCCAAGACT |
| S3_Linker1_1705_for | GTCAGTCAGTCAGTCAGTCAATGTTCTTGTATGTGAGAAC |
| S3_Linker1_1810_for | GTCAGTCAGTCAGTCAGTCAATTGAAGCTGAGTCCTCTGT |
| S3_Linker1_1915_for | GTCAGTCAGTCAGTCAGTCAAGTTCCATTGGGAAGGTCTG |
| S3_Linker1_2020_for | GTCAGTCAGTCAGTCAGTCACTTCTTATCGTTCAGGCTCT |
| S3_Linker2_144_rev | GGATCCAATTGGCCAACTCGAGTGCATATTGCTGCAAATT |
| S3_Linker2_249_rev | GGATCCAATTGGCCAACTCGTCTGTGCTTCAAAAGTGCAT |
| S3_Linker2_354_rev | GGATCCAATTGGCCAACTCGATACAAATCTGGTAGAAACT |
| S3_Linker2_459_rev | GGATCCAATTGGCCAACTCGGATGTGTGTTTCCTCAGATT |
| S3_Linker2_564_rev | GGATCCAATTGGCCAACTCGTTGTCTTATAGTGAATAGTC |
| S3_Linker2_669_rev | GGATCCAATTGGCCAACTCGGGCAAGCTTGCGCATTGTTC |
| S3_Linker2_774_rev | GGATCCAATTGGCCAACTCGGGACATTTGAGACAGCTTGC |
| S3_Linker2_879_rev | GGATCCAATTGGCCAACTCGCATCAGCAGGAATTTGGACC |
| S3_Linker2_984_rev | GGATCCAATTGGCCAACTCGGGGTTCCTTCCATCCAAAGA |
| S3_Linker2_1089_rev | GGATCCAATTGGCCAACTCGTGGAATTTTCTCCTCATTCT |
| S3_Linker2_1194_rev | GGATCCAATTGGCCAACTCGCAAATCACCTACATCTTTAC |
| S3_Linker2_1299_rev | GGATCCAATTGGCCAACTCGGAGCTCTATCCAGCTTGAAT |
| S3_Linker2_1404_rev | GGATCCAATTGGCCAACTCGCATTATGTATTCTGTGGCTC |
| S3_Linker2_1509_rev | GGATCCAATTGGCCAACTCGCCTTCCCTCCTTAGTTCTAC |
| S3_Linker2_1614_rev | GGATCCAATTGGCCAACTCGTGGGTCAGTGAGAGAAAACT |
| S3_Linker2_1719_rev | GGATCCAATTGGCCAACTCGCACATACAAGAACATGGGCC |
| S3_Linker2_1824_rev | GGATCCAATTGGCCAACTCGGGACTCAGCTTCAATCATAC |
| S3_Linker2_1929_rev | GGATCCAATTGGCCAACTCGCTTCCCAATGGAACTTTCCT |
| S3_Linker2_2034_rev | GGATCCAATTGGCCAACTCGCTGAACGATAAGAAGCAGTT |
| S3_Linker2_2139_rev | GGATCCAATTGGCCAACTCGAGAAGCATTAAGCAAAACCC |
| S4_Linker1_33_for | GTCAGTCAGTCAGTCAGTCAATGAAGGCAAACCTACTGGT |
| S4_Linker1_138_for | GTCAGTCAGTCAGTCAGTCAGTACTCGAGAAGAATGTGAC |
| S4_Linker1_243_for | GTCAGTCAGTCAGTCAGTCAAAATGTAACATCGCCGGATG |
| S4_Linker1_348_for | GTCAGTCAGTCAGTCAGTCAATATGTTATCCAGGAGATTT |
| S4_Linker1_453_for | GTCAGTCAGTCAGTCAGTCACCCAACCACAACACAAACAA |
| S4_Linker1_558_for | GTCAGTCAGTCAGTCAGTCACCAAAGCTGAAAAATTCTTA |
| S4_Linker1_663_for | GTCAGTCAGTCAGTCAGTCAAATGAAAATGCTTATGTCTC |
| S4_Linker1_768_for | GTCAGTCAGTCAGTCAGTCATATTACTGGACCTTACTAAA |
| S4_Linker1_873_for | GTCAGTCAGTCAGTCAGTCAGGCATCATCACCTCAAACGC |
| S4_Linker1_978_for | GTCAGTCAGTCAGTCAGTCAACAATAGGAGAGTGCCCAAA |
| S4_Linker1_1083_for | GTCAGTCAGTCAGTCAGTCAGCCGGTTTTATTGAAGGGGG |
| S4_Linker1_1188_for | GTCAGTCAGTCAGTCAGTCACAAAATGCCGTTAACGGGAT |
| S4_Linker1_1293_for | GTCAGTCAGTCAGTCAGTCAATGGAAAATTTAAATAAAAA |
| S4_Linker1_1398_for | GTCAGTCAGTCAGTCAGTCAGACTCAAATGTGAAGAATCT |
| S4_Linker1_1503_for | GTCAGTCAGTCAGTCAGTCAGAATGCATGGAAAGTGTAAG |
| S4_Linker1_1608_for | GTCAGTCAGTCAGTCAGTCAGGGATCTATCAGATTCTGGC |
| S4_Linker2_152_rev | GGATCCAATTGGCCAACTCGATTCTTCTCGAGTACTGTGT |
| S4_Linker2_257_rev | GGATCCAATTGGCCAACTCGGGCGATGTTACATTTCCCCA |
| S4_Linker2_362_rev | GGATCCAATTGGCCAACTCGTCCTGGATAACATATTCCAT |
| S4_Linker2_467_rev | GGATCCAATTGGCCAACTCGTGTGTTGTGGTTGGGCCATG |
| S4_Linker2_572_rev | GGATCCAATTGGCCAACTCGATTTTTCAGCTTTGGGTATG |
| S4_Linker2_677_rev | GGATCCAATTGGCCAACTCGATAAGCATTTTCATTCTGAT |
| S4_Linker2_782_rev | GGATCCAATTGGCCAACTCGTAAGGTCCAGTAATAGTTCA |
| S4_Linker2_887_rev | GGATCCAATTGGCCAACTCGTGAGGTGATGATGCCGGACC |
| S4_Linker2_992_rev | GGATCCAATTGGCCAACTCGGCACTCTCCTATTGTGACTG |
| S4_Linker2_1097_rev | GGATCCAATTGGCCAACTCGTTCAATAAAACCGGCAATGG |
| S4_Linker2_1202_rev | GGATCCAATTGGCCAACTCGGTTAACGGCATTTTGTGTGC |
| S4_Linker2_1307_rev | GGATCCAATTGGCCAACTCGATTTAAATTTTCCATCCTCT |
| S4_Linker2_1412_rev | GGATCCAATTGGCCAACTCGCTTCACATTTGAGTCATGGA |
| S4_Linker2_1517_rev | GGATCCAATTGGCCAACTCGACTTTCCATGCATTCATTGT |
| S4_Linker2_1622_rev | GGATCCAATTGGCCAACTCGAATCTGATAGATCCCCATTG |
| S4_Linker2_1727_rev | GGATCCAATTGGCCAACTCGGCATATTCTGCACTGCAAAG |
| S7_Linker1_26_for | GTCAGTCAGTCAGTCAGTCAATGAGTCTTCTAACCGAGGT |
| S7_Linker1_131_for | GTCAGTCAGTCAGTCAGTCAAACACCGATCTTGAGGTTCT |
| S7_Linker1_236_for | GTCAGTCAGTCAGTCAGTCAGAGCGAGGACTGCAGCATAG |
| S7_Linker1_341_for | GTCAGTCAGTCAGTCAGTCAGAGATAACATTCCATGGGGC |
| S7_Linker1_446_for | GTCAGTCAGTCAGTCAGTCAGAAGTGGCATTTGGCCTGGT |
| S7_Linker1_551_for | GTCAGTCAGTCAGTCAGTCAGAGAACAGAATGGTTTTAGC |
| S7_Linker1_656_for | GTCAGTCAGTCAGTCAGTCACAAATGGTGCAAGCGATGAG |
| S7_Linker1_761_for | GTCAGTCAGTCAGTCAGTCAGTGCAAATGCAACGGTTCAA |
| S7_Linker1_866_for | GTCAGTCAGTCAGTCAGTCATTACCGTCGCTTTAAATACG |
| S7_Linker2_145_rev | GGATCCAATTGGCCAACTCGCTCAAGATCGGTGTTCTTCC |
| S7_Linker2_250_rev | GGATCCAATTGGCCAACTCGCTGCAGTCCTCGCTCACTGG |
| S7_Linker2_355_rev | GGATCCAATTGGCCAACTCGATGGAATGTTATCTCCCTCT |
| S7_Linker2_460_rev | GGATCCAATTGGCCAACTCGGCCAAATGCCACTTCAGTGG |
| S7_Linker2_565_rev | GGATCCAATTGGCCAACTCGAACCATTCTGTTCTCATGCC |
| S7_Linker2_670_rev | GGATCCAATTGGCCAACTCGCGCTTGCACCATTTGTCTAG |
| S7_Linker2_775_rev | GGATCCAATTGGCCAACTCGCCGTTGCATTTGCACCCCCA |
| S7_Linker2_880_rev | GGATCCAATTGGCCAACTCGTTAAAGCGACGGTAAATGCA |
| S7_Linker2_985_rev | GGATCCAATTGGCCAACTCGAAATGACCATCGTCAGCATC |
| S8_Linker1_27_for | GTCAGTCAGTCAGTCAGTCAATGGATCCAAACACTGTGTC |
| S8_Linker1_132_for | GTCAGTCAGTCAGTCAGTCACTTCGCCGAGATCAGAAATC |
| S8_Linker1_237_for | GTCAGTCAGTCAGTCAGTCAGAAGAATCCGATGAGGCACT |
| S8_Linker1_342_for | GTCAGTCAGTCAGTCAGTCAATACCCAAGCAGAAAGTGGC |
| S8_Linker1_447_for | GTCAGTCAGTCAGTCAGTCACTGGAGACTCTAATATTGCT |
| S8_Linker1_552_for | GTCAGTCAGTCAGTCAGTCAAATGCAGTTGGAGTCCTCAT |
| S8_Linker1_657_for | GTCAGTCAGTCAGTCAGTCAAGACCTCCACTCACTCCAAA |
| S8_Linker1_762_for | GTCAGTCAGTCAGTCAGTCAGATAACAGAGAATAGTTTTG |
| S8_Linker2_146_rev | GGATCCAATTGGCCAACTCGCTGATCTCGGCGAAGCCGAT |
| S8_Linker2_251_rev | GGATCCAATTGGCCAACTCGCTCATCGGATTCTTCTTTCA |
| S8_Linker2_356_rev | GGATCCAATTGGCCAACTCGTTTCTGCTTGGGTATGAGCA |
| S8_Linker2_461_rev | GGATCCAATTGGCCAACTCGTATTAGAGTCTCCAGCCGGT |
| S8_Linker2_566_rev | GGATCCAATTGGCCAACTCGGACTCCAACTGCATTTTTGA |
| S8_Linker2_671_rev | GGATCCAATTGGCCAACTCGAGTGAGTGGAGGTCTCCCAT |
| S8_Linker2_776_rev | GGATCCAATTGGCCAACTCGCTATTCTCTGTTATCTTCAG |
| S8_Linker2_881_rev | GGATCCAATTGGCCAACTCGAAGGGTGTTTTTTATTACTA |
| Seg6 Linker1_21 for | GTCAGTCAGTCAGTCAGTCAATGAATCCAAATCAGAAAA |
| Seg6 Linker1_126 for | GTCAGTCAGTCAGTCAGTCACATTCAATTCAAACTGGAAG |
| Seg6 Linker1_ 231 for | GTCAGTCAGTCAGTCAGTCAACCGGCAATTCATCTCTTTG |
| Seg6 Linker1_ 336 for | GTCAGTCAGTCAGTCAGTCATTTATTTCATGTTCTCACTT |
| Seg6 Linker1_441 for | GTCAGTCAGTCAGTCAGTCAAGGGCCTTAATGAGCTGCCC |
| Seg6 Linker1_546 for | GTCAGTCAGTCAGTCAGTCACTAACAATCGGAATTTCAGG |
| Seg6 Linker1_651 for | GTCAGTCAGTCAGTCAGTCAACACAAGAGTCTGAATGTGC |
| Seg6 Linker1_756 for | GTCAGTCAGTCAGTCAGTCAGGGAAGGTTACTAAATCAAT |
| Seg6 Linker1_861 for | GTCAGTCAGTCAGTCAGTCATGGCATGGTTCGAACCGGCC |
| Seg6 Linker1_966 for | GTCAGTCAGTCAGTCAGTCAGGAACAGGCAGCTGTGGTCC |
| Seg6 Linker1_1071 for | GTCAGTCAGTCAGTCAGTCAAGTTCCAGACATGGGTTTGA |
| Seg6 Linker1_1176 for | GTCAGTCAGTCAGTCAGTCAGGGTATAGCGGGAGTTTCGT |
| Seg6 Linker1_1281 for | GTCAGTCAGTCAGTCAGTCAATCTGGACTAGTGCGAGCAG |
| Seg6 Linker2_140 rev | GGATCCAATTGGCCAACTCGAGTTTGAATTGAATGGCTAA |
| Seg6 Linker2_245 rev | GGATCCAATTGGCCAACTCGAGATGAATTGCCGGTTAATA |
| Seg6 Linker2_350 rev | GGATCCAATTGGCCAACTCGAGAACATGAAATAAAGGGCT |
| Seg6 Linker2_455 rev | GGATCCAATTGGCCAACTCGGCTCATTAAGGCCCTATAAG |
| Seg6 Linker2_560 rev | GGATCCAATTGGCCAACTCGAATTCCGATTGTTAGCCAGC |
| Seg6 Linker2_665 rev | GGATCCAATTGGCCAACTCGTTCAGACTCTTGTGTCCTCA |
| Seg6 Linker2_770 rev | GGATCCAATTGGCCAACTCGTTTAGTAACCTTCCCCTTTT |
| Seg6 Linker2_875 rev | GGATCCAATTGGCCAACTCGGTTCGAACCATGCCAATTGT |
| Seg6 Linker2_980 rev | GGATCCAATTGGCCAACTCGACAGCTGCCTGTTCCATCTT |
| Seg6 Linker2_1085 rev | GGATCCAATTGGCCAACTCGCCCATGTCTGGAACTGTGAC |
| Seg6 Linker2_1190 rev | GGATCCAATTGGCCAACTCGACTCCCGCTATACCCTGACC |
| Seg6 Linker2_1295 rev | GGATCCAATTGGCCAACTCGCGCACTAGTCCAGATTGTTT |
| Seg6 Linker2_ 1382rev | GGATCCAATTGGCCAACTCGCTTGTCAATGGTGAATGGCA |
| Seg5 Linker1_46 for | GTCAGTCAGTCAGTCAGTCAATGGCGTCTCAAGGCACCAA |
| Seg5 Linker1_151 for | GTCAGTCAGTCAGTCAGTCAATTGGACGATTCTACATCCA |
| Seg5 Linker1_256 for | GTCAGTCAGTCAGTCAGTCATTTGACGAAAGGAGAAATAA |
| Seg5 Linker1_361 for | GTCAGTCAGTCAGTCAGTCAAGAGAACTCATCCTTTATGA |
| Seg5 Linker1_466 for | GTCAGTCAGTCAGTCAGTCATCCAATTTGAATGATGCAAC |
| Seg5 Linker1_571 for | GTCAGTCAGTCAGTCAGTCATCTGGAGCCGCAGGTGCTGC |
| Seg5 Linker1_676 for | GTCAGTCAGTCAGTCAGTCAAATGGACGAAAAACAAGAAT |
| Seg5 Linker1_781 for | GTCAGTCAGTCAGTCAGTCACGGAACCCAGGGAATGCTGA |
| Seg5 Linker1_886 for | GTCAGTCAGTCAGTCAGTCATATGGACCTGCCGTAGCCAG |
| Seg5 Linker1_991 for | GTCAGTCAGTCAGTCAGTCAATCAGACCAAATGAGAATCC |
| Seg5 Linker1_1096 for | GTCAGTCAGTCAGTCAGTCAAAGGTGCTCCCAAGAGGGAA |
| Seg5 Linker1_1201 for | GTCAGTCAGTCAGTCAGTCATGGGCCATAAGGACCAGAAG |
| Seg5 Linker1_1306 for | GTCAGTCAGTCAGTCAGTCAGACAGAACAACCATTATGGC |
| Seg5 Linker1_1411 for | GTCAGTCAGTCAGTCAGTCAGTGTCTTTCCAGGGGCGGGG |
| Seg5 Linker2_165 rev | GGATCCAATTGGCCAACTCGGTAGAATCGTCCAATTCCAC |
| Seg5 Linker2_270 rev | GGATCCAATTGGCCAACTCGTCTCCTTTCGTCAAAAGCAG |
| Seg5 Linker2_375 rev | GGATCCAATTGGCCAACTCGAAGGATGAGTTCTCTCATCC |
| Seg5 Linker2_480 rev | GGATCCAATTGGCCAACTCGATCATTCAAATTGGAATGCC |
| Seg5 Linker2_585 rev | GGATCCAATTGGCCAACTCGACCTGCGGCTCCAGACCTCC |
| Seg5 Linker2_690 rev | GGATCCAATTGGCCAACTCGTGTTTTTCGTCCATTCTCAC |
| Seg5 Linker2_795 rev | GGATCCAATTGGCCAACTCGATTCCCTGGGTTCCGGCTCT |
| Seg5 Linker2_900 rev | GGATCCAATTGGCCAACTCGTACGGCAGGTCCATACACAC |
| Seg5 Linker2_1005 rev | GGATCCAATTGGCCAACTCGCTCATTTGGTCTGATTAGGC |
| Seg5 Linker2_1110 rev | GGATCCAATTGGCCAACTCGTCTTGGGAGCACCTTCGTCC |
| Seg5 Linker2_1215 rev | GGATCCAATTGGCCAACTCGGGTCCTTATGGCCCAGTACC |
| Seg5 Linker2_1320 rev | GGATCCAATTGGCCAACTCGAATGGTTGTTCTGTCAAAAG |
| Seg5 Linker2_1425 rev | GGATCCAATTGGCCAACTCGCCCCTGGAAAGACACATCTT |
| Seg5 Linker2_1530 rev | GGATCCAATTGGCCAACTCGCTCCTCTGCATTGTCTCCGA |
